# Mountain lion genomes provide insights into genetic rescue of inbred populations

**DOI:** 10.1101/482315

**Authors:** Nedda F. Saremi, Megan A. Supple, Ashley Byrne, James A. Cahill, Luiz Lehmann Coutinho, Love Dalén, Henrique V. Figueiró, Warren E. Johnson, Heather J. Milne, Stephen J. O’Brien, Brendan O’Connell, David P. Onorato, Seth P.D. Riley, Jeff A. Sikich, Daniel R. Stahler, Priscilla Marqui Schmidt Villela, Christopher Vollmers, Robert K. Wayne, Eduardo Eizirik, Russell B. Corbett-Detig, Richard E. Green, Christopher C. Wilmers, Beth Shapiro

**Author notes:** Contributed equally to this work (Nedda F. Saremi, Megan A. Supple).

## Abstract

Across the geographic range of mountain lions, which includes much of North and South America, populations have become increasingly isolated due to human persecution and habitat loss. To explore the genomic consequences of these processes, we assembled a high-quality mountain lion genome and analyzed a panel of resequenced individuals from across their geographic range. We found strong geographical structure and signatures of severe inbreeding in all North American populations. Tracts of homozygosity were rarely shared among populations, suggesting that assisted gene flow would restore local genetic diversity. However, the genome of an admixed Florida panther that descended from a translocated individual from Central America had surprisingly long tracts of homozygosity, indicating that genomic gains from translocation were quickly lost by local inbreeding. Thus, to sustain diversity, genetic rescue will need to occur at regular intervals, through repeated translocations or restoring landscape connectivity. Mountain lions provide a rare opportunity to examine the potential to restore diversity through genetic rescue, and to observe the long-term effects of translocation. Our methods and results provide a framework for genome-wide analyses that can be applied to the management of small and isolated populations.

## Main

The ancestors of the mountain lion, also known as the puma or cougar, colonized North America approximately 6 million years ago (mya)^1^–^3^. Although their Pliocene fossil record is sparse, previous mitochondrial analyses suggested that mountain lions diverged from the extinct North American cheetah-like cat *Miracinonyx* around 3.2 mya^4^. The geographic origins of the species remain contested; the oldest unequivocal mountain lion fossil dates to 1.2 to 0.8 mya from Argentina^5^, indicating that the ancestral mountain lion lineage eventually dispersed southward following the re-emergence of the Panamanian isthmus ∼2.5 mya^6,7^. In North America, the oldest mountain lion fossils to date are found at sites across the continent of Rancholabrean age^8^, which provides a current maximum age of ∼200 thousand years ago (kya)^9^. Previous analyses of mitochondrial and microsatellite data estimated a common ancestor of all living North American mountain lions within the last 20,000 years^10^. These data also indicate that today’s puma genetic diversity traces its origins to Eastern South America^10^. Together, the North American fossil record and genetic data have been interpreted as reflecting a North American origin of the species, a local extinction in North America sometime during the late Pleistocene, and recolonization from South America as the climate warmed after the last ice age^10,11^.

Today, mountain lions are among the most widely-distributed mammals in the Western hemisphere, ranging from Canada’s Yukon to the southern tip of South America (Fig. 1)^12^–^14^. During the 19th and 20th centuries, bounty hunting reduced, and in some cases extirpated, mountain lion populations across North America^15^, restricting them to the North American West and the southern tip of Florida. By the middle of the 20th century, hunting quotas and some outright bans^13^ allowed mountain lion populations to increase and recolonize parts of their former range. Human development, however, has continued to degrade and fragment much of their habitat. Although some mountain lion populations today are large and well-connected^16^, others are small and fragmented (e.g. Santa Ana, CA^17^), and/or critically endangered (e.g. Florida^18^). Many populations are experiencing increased isolation with the expansion of highways, residential development and agriculture^17,19^. Loss of diversity may be a common situation for top predators, as their population densities are usually low. Consequently, even moderate levels of fragmentation will greatly affect their genomic diversity.

**Figure 1:**
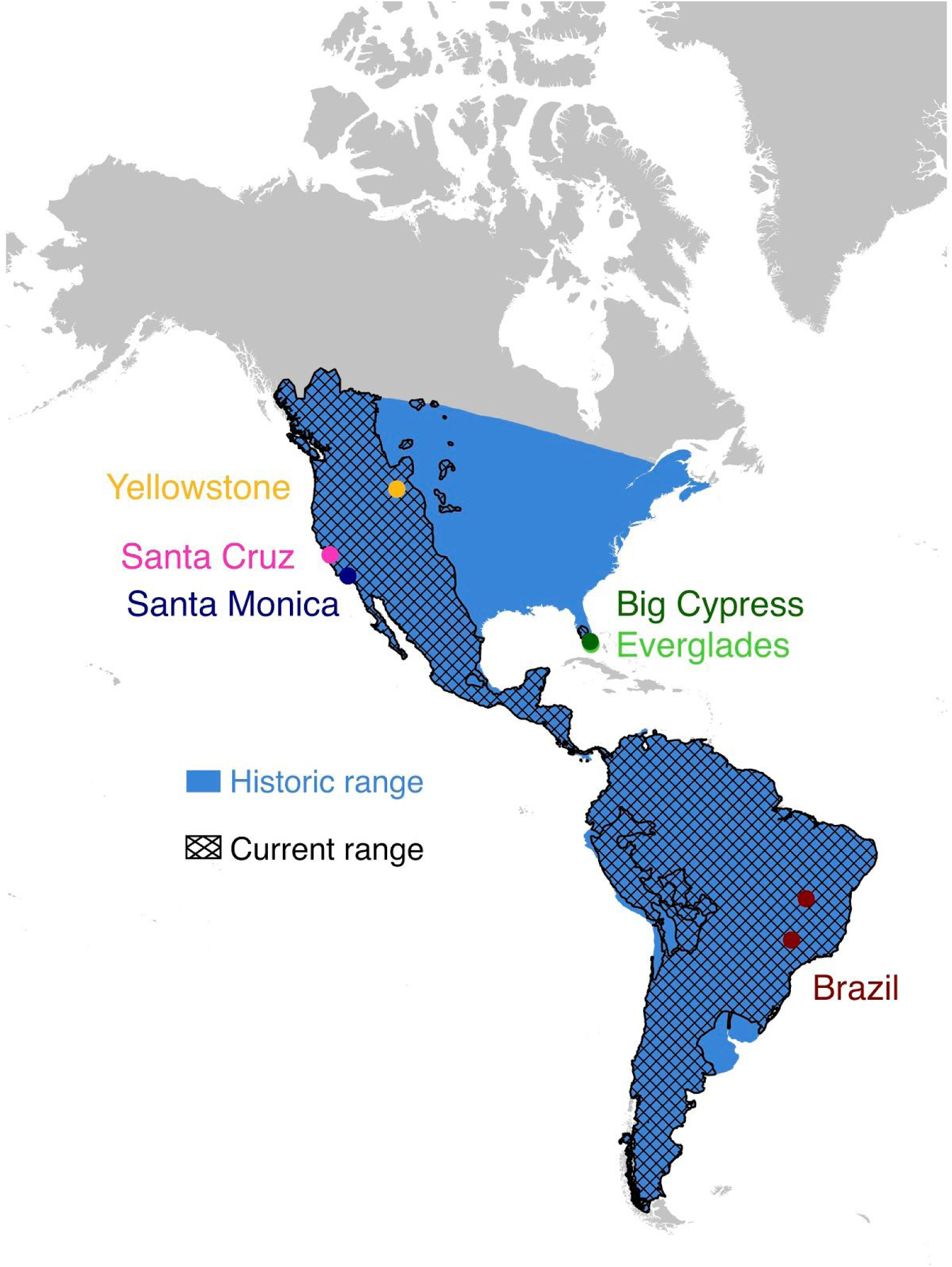
Mountain lion range past and present. The current range of mountain lions (hashed) compared to their historic range (blue). Circles denote the geographic coordinates of the mountain lion populations sampled in this study. Current range data is from the IUCN Red List of Threatened Species^90^. Historic range data is approximated based on prior reports^12,13^.

The consequences of human development on mountain lion genetic diversity and fitness have been well documented, particularly in Florida, where they are a federally protected subspecies commonly called the Florida panther. Most of the breeding population persists in the interior parts of southern Florida in large tracts of public lands and fragmented tracts of private land. By the 1990s, the Florida panther population was suffering from reproductive failure and phenotypic defects associated with inbreeding^18,20^. To rescue the panther population from extinction, eight female Texas mountain lions were released in South Florida in 1995, of which five successfully produced offspring. After one generation, the occurrence of phenotypic defects among Florida panthers was reduced considerably and reproduction was restored^21,22^. Today, all Florida panthers share common ancestry with the five translocated females.

Florida panthers in Everglades National Park are partially isolated from the core population that persisted in Big Cypress National Preserve by a semipermeable barrier associated with hydrologic fluctuations of the Everglades. Intriguingly, during the 1990s, the Everglades panthers did not show a high incidence of inbreeding-associated phenotypes, as was seen in the Big Cypress population. This may have been due to poorly documented attempts at population augmentation during the 1950s and 1960s, when captive-bred Florida panthers with mixed Central American ancestry were introduced into Everglades National Park. The ancestry of the introduced individuals was unclear at the time of release, though it was known that the Everglades population had greater reproductive success than wild Florida panthers. The true ancestry, and reason for greater reproductive success of the captive population, was later determined through genotyping^23^.

Here, we reconstruct the last two million years of mountain lion demographic history by generating and analyzing a high-quality, chromosome-scale assembly from an individual sampled in the Santa Cruz Mountains (northern California, USA), along with nine resequenced genomes from mountain lions from North and South America. We estimate the timing of divergence between North and South American mountain lions and describe how human-caused fragmentation of their habitats has impacted their connectivity and genomic diversity. We use shared tracts of homozygosity to predict the effectiveness of assisted gene flow in restoring lost genetic diversity. Finally, we analyzed the genome of a Florida panther with admixed ancestry that was collected 30 years after the first release of Central American admixed mountain lions into the Everglades. This genome allowed us to assess the long-term efficacy of inter-population admixture as a means to rescue mountain lions from the deleterious effects of inbreeding.

## Results

### Genome Assembly and Variant Calling

We assembled a *de novo* nuclear genome for a wild male mountain lion (SC36) from the Santa Cruz Mountains using a combination of shotgun Illumina, long-range linking Illumina, and Oxford Nanopore Technology (ONT) data^24^ (see Methods). Our PumCon1.0 assembly has a BUSCO^25^ score of 93.4%, a scaffold N50 of 100 Mb, and 87.6% of the genome represented on 26 autosomal scaffolds, each larger than 20 Mb (Supplementary Tables 1,2). Although our ONT coverage was only 1.2X, the use of these data for gap-filling recovered an additional 5.74% of the genome sequence, which we error-corrected by re-mapping the Illumina reads (Supplementary Table 1).

We obtained whole-genome resequencing data from nine additional mountain lions from seven locations in North and South America (Fig. 1 and Supplementary Table 3), and aligned the data to our reference assembly (PumCon1.0) for variant calling. We filtered variants to produce a final call set of 8 million variable sites, which decreased to 166,037 variable sites after filtering for linkage disequilibrium (see Methods).

### Demographic History

We first reconstructed mountain lion demographic history using both mitochondrial and nuclear genomes. We estimate that the most recent common maternal ancestor of all sampled mountain lions lived ∼300 kya (Fig. 2A), and that North American mitochondrial haplotypes cluster together, sharing an inferred common maternal ancestor 31-11 kya. The North American clade excludes the Florida Everglades mountain lion (EVG21), which has a mitochondrial ancestry that is distinct from the rest of North America, consistent with the previously reported mixed ancestry of this individual^18^. Pairwise sequentially Markovian coalescent (PSMC) modelling of the nuclear genomic data also suggests that two mountain lion lineages, one represented by the two Brazilian individuals and the other representing all individuals sampled in North America, diverged around 300 kya (Fig. 2B and Supplementary Fig. 4). The North American mountain lion population size increased after this divergence, presumably as they expanded into unoccupied habitats in Central and North America. Populations on both continents were largest around 130 kya, during the warmest part of Marine Isotope Stage (MIS) 5, and then declined throughout the MIS 4-2 cold stages, reaching their current small sizes by the peak of the last ice age 25-20 kya. The spike in effective population size observed for EVG21 likely does not reflect true demographic history, but is instead consistent with mixed ancestry comprising two divergent lineages^26^.

**Figure 2:**
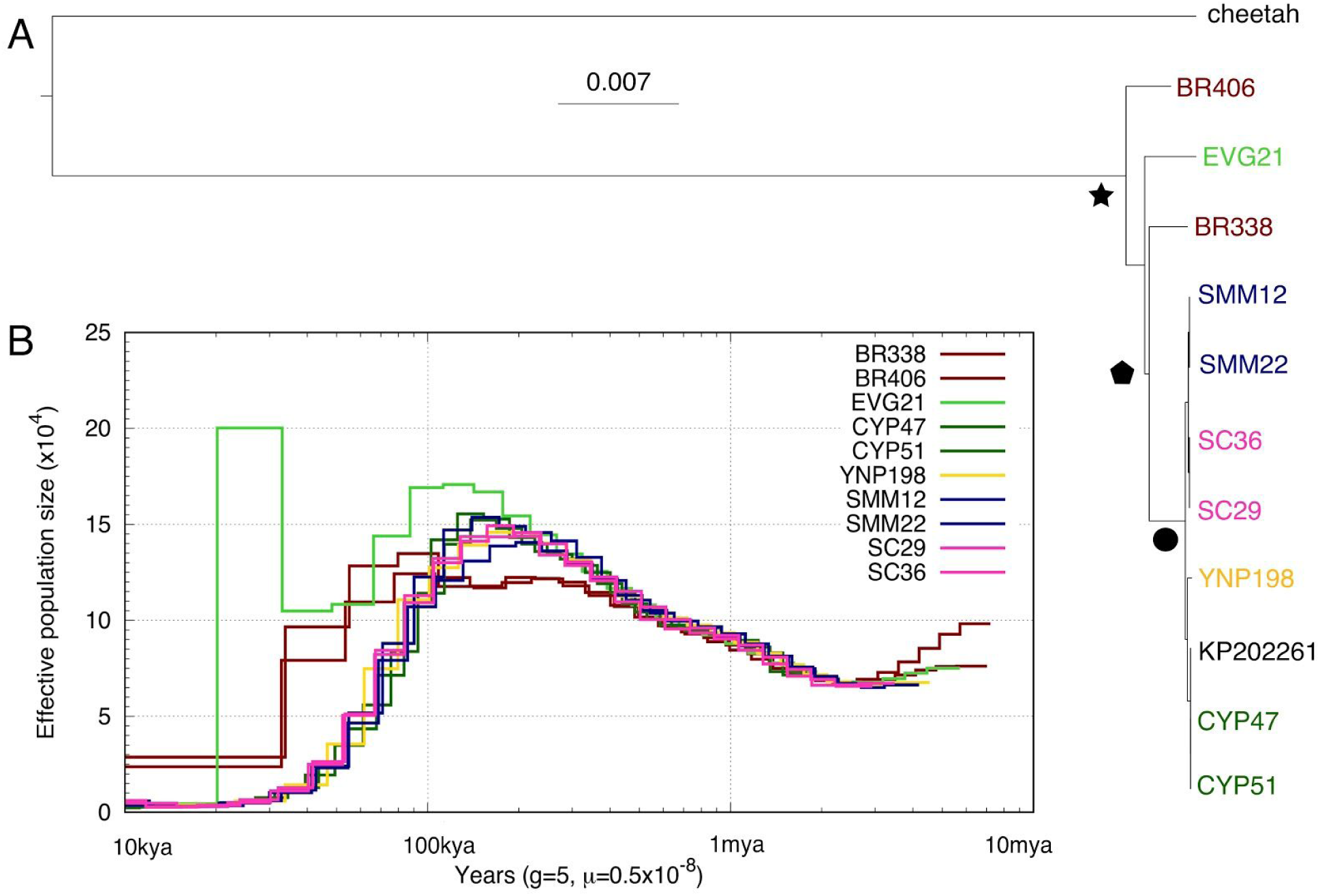
Demographic history of mountain lions. **A)** Mitochondrial maximum likelihood phylogeny of the ten mountain lions in this study plus a mountain lion from Big Cypress (KP202261.1) and the African cheetah (KP202271.1) as the outgroup. Using a mitochondrial divergence rate of 1.15%/Myr^10,70^, we estimate a common maternal ancestor of all mountain lions 278,000 ± 5,639 years ago (star; 100% bootstrap support), divergence between North American and South American mitochondrial lineages 201,000 ± 1,952 years ago (pentagon; 63% bootstrap support), and a common maternal ancestor of North American all mountain lions 21,000 ± 10,412 years ago (circle; 100% bootstrap support). **B)** Inferred changes in effective population size (Ne) over time using the pairwise sequentially Markovian coalescent (PSMC) model^78^ for the ten mountain lions. We assume a generation time of 5 years and a per generation mutation rate of 0.5e-8^79^. The demographic histories of our North (green, blue, yellow, pink lines) and South (red lines) American samples diverge ∼300,000 years ago. The PSMC plot for EVG21 shows an increase in inferred Ne within the 110,000 that is probably attributable to its hybrid ancestry.

### Population Structure

The nuclear genomic data was subsequently used to characterize geographic structure among mountain lion populations (Fig. 3). We performed principal component analysis (PCA) of 166,037 overlapping SNPs and found strong evidence of geographic structure. Populations represented by more than one individual clustered together and 52% of the genetic variance within and among populations was explained by the first two principal components (Fig. 3A). These PCs divide North and South America, revealing a gradient of relatedness from east to west across North America, and separate EVG21 from the two Florida panthers from the Big Cypress population. We estimated a consensus nuclear phylogeny and found further evidence of geographic structure, with the highest likelihood tree including a single migration event from a South (or Central) American lineage into the Everglades lineage (Fig. 3B and Supplementary Fig. 6,7). Finally, our cluster assignment tests also partitioned the data geographically, with the two continents separating at K=2 and additional groups emerging along the same gradients described in the PCA (Fig. 3C and Supplementary Fig. 8). Notably, EVG21 shares ancestry with both Florida and Brazil with K=3 and K=4.

**Figure 3:**
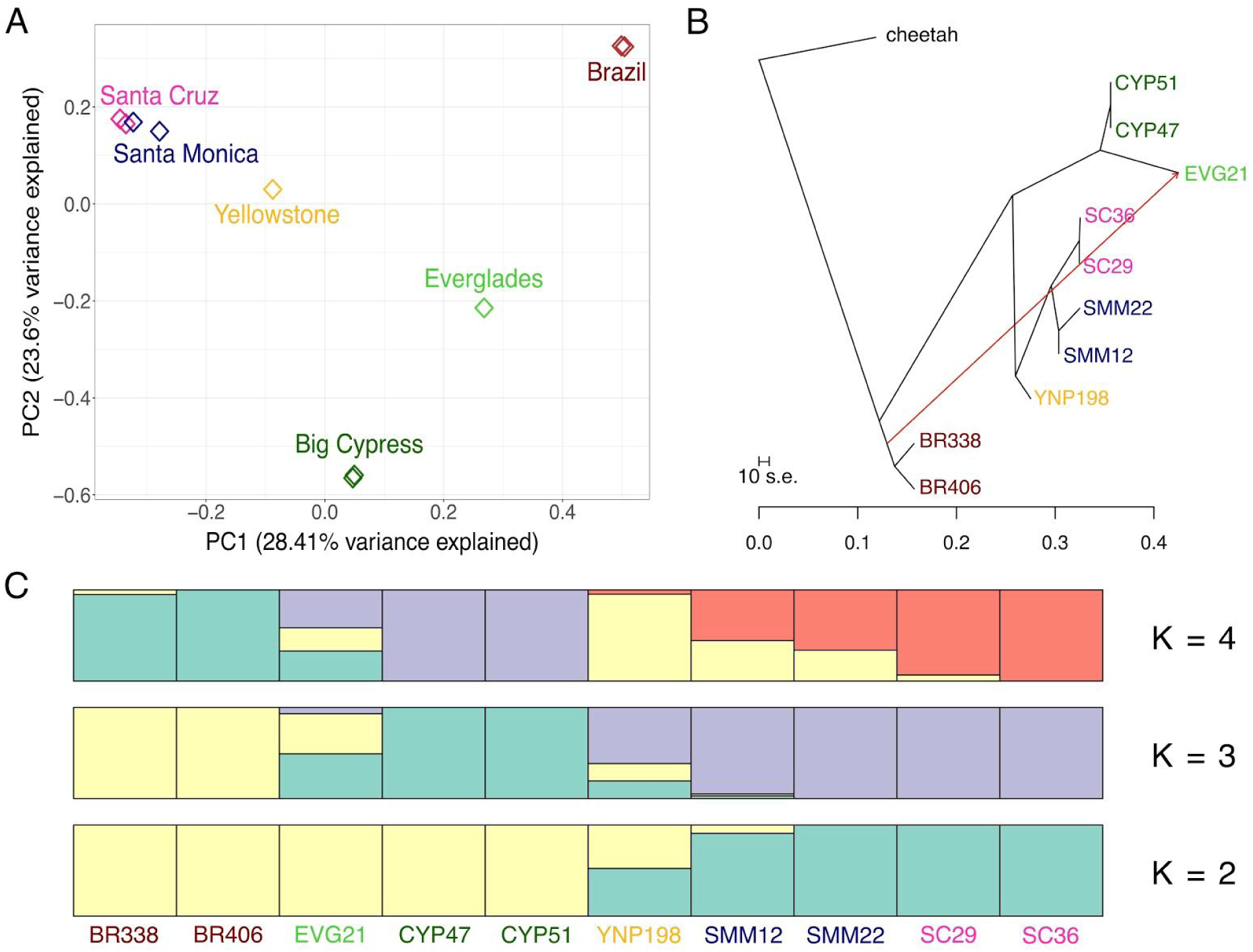
Stratification of mountain lions based on geographic population. **A)** Principal component analysis^80^ of 166,037 sites separates the sampled mountain lions based on geography. The first component primarily separates South and North American mountain lions, while the second component distinguishes the variation within North America. All Californian mountain lions (Santa Cruz and Santa Monica) cluster closely. **B)** TreeMix^82^ analysis, using the cheetah as the outgroup, indicates the best tree separates mountain lions based on geographic population and includes one migration event (weight = 0.453911) from the branch of South American diversity into the admixed Everglades mountain lion (EVG21). **C)** The mean of 10 permuted matrices of STRUCTURE^83^ analyses for each of K=2 through 4, performed using CLUMPP^85^. K=3 had the largest delta K value (Supplementary Fig. 7)^84^.

### Heterozygosity and Inbreeding

Next, we examined the extent of inbreeding in our mountain lion samples. For each individual, we estimated the average genome-wide heterozygosity and identified runs of homozygosity (ROH) across the 26 largest autosomal scaffolds (Fig. 4, see Methods). The distribution of ROH across the genome varied among scaffolds and individuals (Fig. 4A and Supplementary Fig. 10), as did average genome-wide heterozygosity and proportion of the genome in ROH (Fig. 4B). The two mountain lions from Brazil were the least inbred, with the highest levels of heterozygosity and smallest proportions of their genomes in ROH. Conversely, the Big Cypress Florida panthers sampled prior to the 1995 genetic rescue were the most inbred, with the lowest levels of heterozygosity and the largest proportions of their genomes in ROH, consistent with the phenotypic defects recorded in these individuals^18^. The other North American mountain lions fell between these two extremes, with all individuals showing some amount of close inbreeding. Of the two individuals from the Santa Monica Mountains, SMM12 appeared to be less inbred than SMM22, with higher heterozygosity and a lower proportion of its genome in ROH. This is consistent with their origins, as genetic analysis suggests that SMM22 was likely born in the small and more isolated Santa Monica Mountains population south of US freeway 101, whereas SMM12 was first observed in the larger and more connected population north of US 101 and dispersed into the Santa Monica Mountains as a subadult^19^.

**Figure 4:**
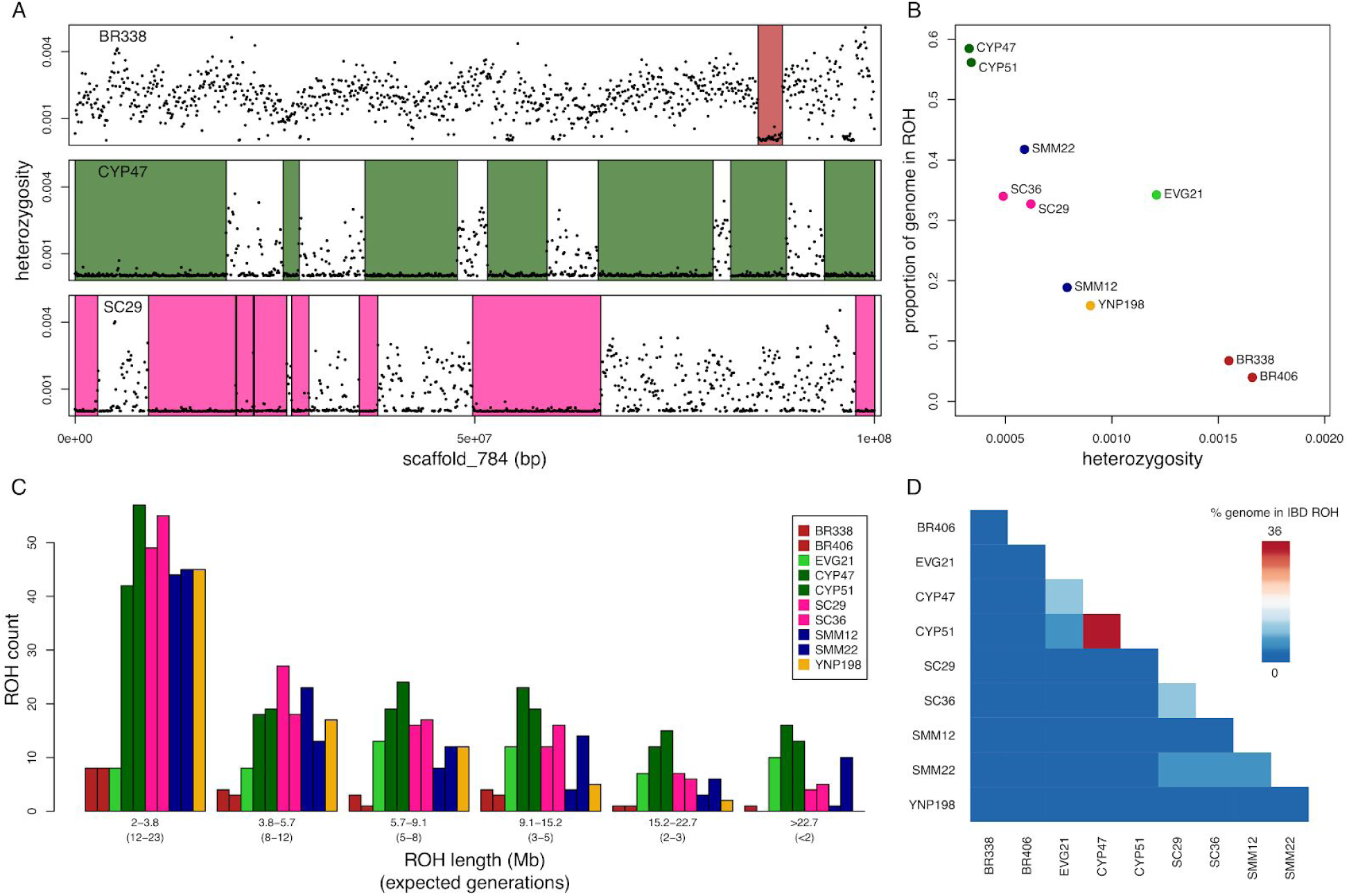
Heterozygosity and Runs of Homozygosity. **A)** Sliding window heterozygosity (black dots) and called ROH (colored boxes) across a single scaffold for three mountain lions from three different populations (Brazil, Big Cypress, and Santa Cruz). Plots for all mountain lions are provided as Supplemental Fig. 9. **B)** Average genome wide heterozygosity versus the proportion of the genome in ROH for the ten mountain lions sequenced. **C)** Distribution of lengths of ROH. The length in Mb is indicated, as is the associated expected number of generations since the individual’s maternal and paternal lineages shared a common ancestor. **D)** Heat map showing the percent of the genome that are ROH that are shared IBD between pairs of mountain lions.

EVG21, the admixed Florida panther from the Everglades population, was an outlier in the general correlation between heterozygosity and proportion of the genome in ROH. The proportion of EVG21’s genome in ROH was high relative to the expectation based on its average genome wide heterozygosity. This is consistent with both ancestral admixture resulting in a more diverse genetic background and close inbreeding leading to long tracts of homozygosity.

To better explore inbreeding history, we examined the distribution of ROH tract lengths in each mountain lion and correlated those lengths with the expected number of generations since its maternal and paternal lineages shared a common ancestor (Fig. 4C, see Methods). Long ROH (>15.2 Mb) occur due to close inbreeding (a common ancestor <3 generations ago). Short ROH (<5.7 Mb) occur due to shared ancestors further back in time (> 8 generations ago). The Big Cypress panthers each had many long ROH, which we estimated to reflect common ancestors within the last three generations. The individuals from Santa Cruz and the Santa Monica Mountains had many intermediate and long ROH (> 5.7 Mb), suggesting that these populations also experienced close inbreeding. The admixed Everglades panther, EVG21, had a small number of short ROH (<5.7 Mb), similar to the Brazilian mountain lions, but mostly long ROH (>15.2 Mb), similar to the more inbred Florida individuals. This combination can be attributed to EVG21’s complex history of admixture and inbreeding. EVG21 is both outbred and an offspring of an inbreeding event—the sire of EVG21 was also EVG21’s half brother^18^ (Supplementary Fig. 3). The peak of the ROH length distribution for EVG21 occurs at 5.7-9.1 Mb, indicating that EVG21’s maternal and paternal lineages possibly shared a common ancestor as far back as 5-8 generations, shortly after the admixture event that occurred 6-9 generations prior^18^.

Although North American mountain lions all have long ROH, these tracts were generally not identical by descent (IBD) between individuals (Fig. 4D). Long ROH that are shared IBD between individuals are concerning because they represent regions of the genome with no genetic diversity in the four haplotypes analyzed. Of the mountain lions sequenced, only the two individuals from Big Cypress (CYP47 and CYP51) shared a considerable proportion (36%) of their genomes in ROH that are IBD. The mountain lions from Santa Cruz (SC29 and SC36) shared 12% of their genomes in IBD ROH, whereas the mountain lions that originate from different areas in and near the Santa Monica Mountains (SMM12 and SMM22) shared only 4%. Individuals from Santa Cruz and the Santa Monica Mountains shared between 3% and 5%. This result suggests that distinct inbreeding histories underlie the genealogy of mountain lions in different populations; while all North American populations show signs of inbreeding, different populations are fixed for different variants and considerable genetic variation still exists when considering the species as a whole.

## Discussion

We present a high-quality, chromosome-scale assembly of a mountain lion genome. Our genome allows us to reconstruct the demographic history of the species as well as to measure genome-wide heterozygosity and ROH, which is less practical with lower-quality or reference-guided genome assemblies. Our assembly strategy combined short-read Illumina data with long-read data from Oxford Nanopore Technologies to generate a scaffold N50 of 100 Mb, making this one of the most contiguous genomes assembled to date. Although our coverage from ONT was only 1.2X, the addition of these data to fill gaps between HiC-linked scaffolds recovered over 140,000,000 bp of sequence, which we error-corrected by re-mapping the lower-error Illumina reads. This approach presents a less costly alternative to recovering missing sequence than alternate strategies that begin, for example, with high-coverage long-read data^27^.

Our analyses of ten complete mountain lion genomes revealed the dynamic history of a very successful species whose population size is now reduced across much of its range. We show that extant North American mountain lions are descended from a population that dispersed from South America during the Late Pleistocene ∼300 kya. Previously, estimates of the timing of divergence between North and South American populations based on rapidly evolving microsatellites and partial mitochondrial genomes, combined with the fossil record, led to the hypothesis of a North American origin of the species, followed by a Pleistocene local extinction in North America and a recolonization from South America within the last 20,000 years^10,11^. Our estimates using complete nuclear and mitochondrial genomes, in combination with a recently identified mountain lion fossil in South America dating to 1.2-0.8 mya^5^, support a different model in which mountain lions originated as a species in South America, colonized Central and North America around 300 kya, and persisted in these areas as a distinct population until the present day. A single-colonization model is also consistent with the fossil record, in which the oldest mountain lion fossils are from South America and the oldest North American fossils date to around the time that our North American nuclear genomic lineages diverged from the two lineages from Brazil^8,9^.

North American mountain lions were successful throughout MIS 5, a warm interglacial period during which forests were prevalent across much of the continent^28^. As the climate cooled into the MIS 4 ice age, forests were replaced by more open and exposed grasslands, and megafaunal prey decreased in abundance^29^. Mountain lion populations declined such that all individuals sampled in North America shared a common maternal ancestor that lived around the peak of the last ice age some 25-20 kya, indicating that mountain lion populations were already small by that time. Intriguingly, mountain lions in South America are known to occupy grassland habitats, for example in Patagonia^15^. Differences in habitat selection between the two continents may reflect competition with distinct sets of carnivoran species. Today, jaguars outcompete mountain lions in moist forest biomes of South America^30^, while wolves outcompete mountain lions in North American grasslands^31^. Furthermore, present differences in habitat selection probably reflect a long history of competition with a much more diverse carnivore guild on both continents during the Pleistocene.

The recent history of mountain lions is marked by human encroachment on their habitat. Even after the abatement of hunting pressure in the mid-20th century, human infrastructure including cities, highways, and agriculture continue to isolate mountain lion populations. Our results highlight the genomic consequences of this isolation, as all North American mountain lion populations are strongly geographically clustered and show signs of close inbreeding.

Florida panthers are among the most well-studied populations with regard to the phenotypic manifestations of isolation and inbreeding. The 1995 introduction of mountain lions from Texas, the geographically most proximate population to Florida panthers, is widely regarded as a successful genetic rescue via translocation. However, Florida panther genetic diversity in the Everglades population had been bolstered several decades earlier, when seven individuals were released into Everglades National Park from a captive breeding facility where mountain lions from Central America had been included in the breeding population. One Florida panther that we sequenced, EVG21, is descended from an admixed individual with Central American ancestry. Her genome is a combination of regions with comparatively high heterozygosity, similar to that observed in the Brazilian mountain lions, and long ROH, similar to the highly inbred Florida panthers. The distribution of the lengths of ROH suggest that her maternal and paternal lineages share a common ancestor that lived shortly after the release of the admixed mountain lions into the Everglades population. This suggests that the genomic consequences of inbreeding happen quickly, with much of the gains from the genetic rescue being erased in only a few generations. EVG21’s genome provides evidence that a consistent effort is required to maintain the benefits of translocation.

The Florida panther population is, however, not the only mountain lion population showing a genetic signature of close inbreeding. Mountain lions in the Santa Monica Mountains and the Santa Cruz Mountains all have long ROH, indicating close inbreeding. Both of these populations are largely isolated from other nearby populations by geography and human development^19,32,33^. In contrast, the individual sampled from Yellowstone National Park had the highest heterozygosity in our panel of North American mountain lions. Although mountain lions in the Yellowstone area were hunted to low densities by the mid 20th century^15^, connectivity between Yellowstone and surrounding wildlands may have facilitated the recovery and maintenance of genetic diversity once hunting pressure was reduced.

In many areas of the current mountain lion range, human land use has reduced the connectivity that is critical to recovery and maintenance of healthy populations. Despite these barriers, gene flow among neighboring populations can be facilitated by enhancing landscape connectivity through coordinated land use planning and by adding bridges or underpasses across freeways. Although mountain lions are capable of traveling long distances, large roads are a major barrier to their movement^17,34^. A model of population dynamics in the Santa Monica Mountains that incorporated landscape connectivity and its effects on genetic diversity predicted a high probability of extinction (99.7%) within 50 years after survival rates decrease due to inbreeding, unless connectivity was increased^35^. Our genomic analyses of the samples from the Santa Monica Mountains also support the effectiveness of population connectivity. The two mountain lions sequenced from the region (SMM12 and SMM22) both reside in the small subpopulation south of US 101 freeway, but SMM12 migrated into the subpopulation from north of US 101^19^. Migrations between these two subpopulations are now rare, but they were probably part of a larger panmictic population prior to the existence of US 101. The genomic analysis of ROH in SMM12 and SMM22 shows that only 4% of their genomes are in ROH that are IBD. This indicates that while both individuals have reduced diversity in a considerable proportion of their genomes, these regions are generally not shared between them. Thus, reconnecting the populations on either side of US 101, as currently proposed via a wildlife crossing over the freeway^36^, would help restore the lost genetic diversity.

Genome-scale data sets are often touted for their utility in conservation planning. However, our results highlight the importance of selecting appropriate metrics when interpreting genomic data. Measures of average genome-wide heterozygosity are the most commonly used metrics to characterize the genetic health of a species, as estimates are relatively simple to generate and are easily comparable among organisms. However, heterozygosity provides only a narrow insight into the health and genetic potential of a species^37^. While in some species average genome-wide heterozygosity is highly correlated with the level of inbreeding estimated using ROH^38^, in systems with admixture, average heterozygosity estimates can be deceptive, as demonstrated with our admixed Everglades mountain lion (EVG21). In this individual, average genome-wide heterozygosity is not indicative of the level of inbreeding. The heterozygosity of EVG21 is almost as high as the Brazilian mountain lions, but EVG21 has a large portion of her genome in ROH. We would infer two very different genetic conditions when considering each metric separately, and thus both heterozygosity and proportion of the genome in ROH should be considered in assessing genomic health. Finally, knowledge of shared ROH, which is not revealed in estimates of average heterozygosity, is critical when designing mitigation plans, as it predicts whether enhancing connectivity would restore lost genetic diversity and helps identify potential candidates for translocation. In this context, this study can serve as a template for future conservation genomic research targeting species living in increasingly fragmented habitats.

## Methods

### SC36 Sequencing Data

#### Illumina libraries

We obtained blood samples from SC36, a wild male mountain lion who lived in the Santa Cruz Mountains in California, USA. We performed two DNA extractions using the Qiagen DNeasy Blood & Tissue kit following the manufacturer’s protocol for non-nucleated erythrocytes using 100µL of blood. We made four indexed Illumina libraries from these two extractions, following the Meyer Kircher protocol^39^, targeting insert sizes of 350bp and 550bp. We created a third DNA extraction for SC36 using the Qiagen Blood and Cell Culture Mini Kit using 500 µL of blood, following the manufacturer’s protocol up until the spooling step. Instead of spooling, we centrifuged the sample, washed with cold 70% EtOH, and further centrifuged. We removed the supernatant and air dried the pellet. We resuspended the pellet in 50 µL of TE buffer at 55°C for 2 hours. We prepared four indexed Illumina libraries from this extraction following the Meyer Kircher protocol^39^, targeting an insert size of 330bp.

We sent out the eight paired-end shotgun libraries for sequencing at UC Berkeley Vincent J. Coates Genomics Sequencing Laboratory on an Illumina HiSeq 2500 (2×150bp, 2×100bp), at UC Santa Cruz Ancient and Degraded Processing Center on an Illumina MiSeq (v3 chemistry, 2×300bp), and at UC San Diego Institute for Genomic Medicine Genomics Center on an Illumina HiSeq 2500 (2×100bp).

#### CHiCago library

We generated a CHiCago library using blood from SC36 using a previously described method ^40^. We sent the library for sequencing at the UC San Diego Institute for Genomic Medicine Genomics Center on an Illumina HiSeq 2500 (2×125bp).

#### Hi-C library

We generated a Hi-C library for SC36 following previously described methods^41^, modifying the protocol to use 1mL of blood, whereby we immobilized chromatin using SPRI beads. We sent the library for sequencing at UC San Diego Institute for Genomic Medicine Genomics Center on a HiSeq 4000 (2×75bp).

#### Oxford Nanopore libraries

We extracted genomic DNA from a whole blood sample from SC36 with the Qiagen Blood and Cell Culture Mini Kit, using 750 µL of starting material. We followed the protocol exactly, up until the spooling step. Instead of spooling, we centrifuged the sample at 5000g for 15 minutes, washed it with cold 70% EtOH, and centrifuged it again at 5000g for 10 minutes. We removed the supernatant and air-dried the pellet. We resuspended the pellet in 50 µL of TE buffer on a shaker at 22°C overnight. We quantified DNA with a Qubit 2.0 dsDNA HS kit at 71.5 ng/µL (3.58 µg total). We verified the DNA size distribution with a pulse-field gel using a 0.75% agarose TAE gel, run at 75V for 16 hours, with the preset 5-150 kb program on the Pippin Pulse power Supply (version 1.3.2), and estimated the DNA fragments to range in size from 20-25 kb in length.

We performed both a Rapid and a 1D^2^ Sequencing run (Oxford Nanopore, SQK-RAD002 and SQK-LSK308). For the Rapid library preparation, we used an input of 200 ng of DNA, as recommended by the manufacturer. We first treated the high molecular weight DNA with 2.5 µL of fragment repair mix and incubated for 1 minute at 30°C, followed by 1 minute at 75°C. We ligated the repaired DNA with Rapid Adapters using Blunt/TA Ligase (NEB). We quantified the Rapid libraries using a Qubit prior to sequencing. The Rapid sequencing libraries were run on an R9.5 flow cell using the NC_48hr_Sequencing_FLO-MIN107_SQK_RAD002 protocol. For the 1D^2^ libraries, we used the recommended 1µg of high molecular weight DNA. We end-repaired and A-tailed the DNA using NEBNext UltraII End-Repair/dA-tailing mix (NEB). We cleaned up the end-repair product using AMPure XP beads and then ligated 1D^2^ adapters using Blunt/TA Ligase (NEB). We performed a subsequent ligation to add the Barcoded Adapter mix (BAM) using Blunt/TA Ligase (NEB). We quantified the 1D^2^ libraries using a Qubit prior to sequencing. The 1D^2^ libraries were run on an R9.5 flow cell using the NC_48hr_Sequencing_FLO-MIN107_SQK_LSK308 protocol. We base called all Fast5 raw reads generated from both sequencing runs using the latest Albacore (Oxford Nanopore proprietary) (version 2.0.2).

### Nuclear Genome Assembly

We removed adapters from the eight Illumina libraries for SC36 with SeqPrep2, using the default parameters except for the quality score cutoff to 15 (-q 15) and minimum length of a trimmed read to 25 bp (-L 25)^42^. We used Trimmomatic^43^ to 1) remove additional Meyer Kircher IS3 adapters^39^ using a seed mismatch of 2 and a simple clip threshold reduced to 5 for the shorter adapter sequence, 2) quality end trim using minimum qualities of 2 or 5 for leading and trailing ends of reads, respectively, 3) window quality trim, using a window size of 4 and a minimum quality of 15, and 4) remove reads shorter than 50 bp. We used this processed shotgun data to assemble a *de novo* genome using the Meraculous-2D Genome Assembler (version 2.2.4), with diploid mode set to 1 and a kmer size of 45^44^. The contig N50 of the shotgun assembly was 17.6 kb and the scaffold N50 was 36.6 kb. We had roughly 45 fold coverage of the genome from the eight Illumina shotgun libraries.

We scaffolded the Meraculous assembled genome using HiRise^40^ (version 2.1.1) run in serial mode using the default parameters with the Chicago and Hi-C libraries as input. The resulting scaffold N50 was 103.8 Mb (Supplementary Fig. 1).

Given that the mountain lion used for the SC36 shotgun assembly was a male, we had difficulty in assembling the X and Y chromosomes with Meraculous run in diploid mode 1. We identified potential X and Y scaffolds in the SC36 HiRise assembly by using Exonerate^45^ (version 2.2.0) to align to known domestic cat X and Y chromosome genes^46^. We identified two scaffolds (scaffolds 2173 and 1964) which had numerous X chromosome mapping genes, and validated these scaffolds as being X chromosome scaffolds based on coverage of Illumina data in SC36. We removed both scaffolds from the HiRise assembly. We identified one scaffold that confidently mapped exclusively to a Y chromosome gene, but were unable to validate it based on coverage, so we did not remove it from our HiRise assembly.

To generate an assembly for the X chromosome, we used shotgun data generated from a female mountain lion from the Santa Monica Mountains, SMM13 (Section Mountain lion resequencing data), using Meraculous^44^ (version 2.2.4) in diploid mode 1 and a kmer size of 45. We determined sex chromosome scaffolds in the SMM13 genome assembly using Exonerate^45^ (version 2.2.0) by identifying scaffolds that align to known domestic cat X genes^46^. We identified 3 large scaffolds in this way (scaffolds X1, X2, and X3), and validated them based on coverage for both SC36 and SMM13. For SMM13, we found that these scaffolds had roughly the same coverage as the average genome wide coverage. For SC36, we found these scaffolds had roughly half coverage of the average genome coverage. We added these three X chromosome scaffolds to the HiRise assembly for SC36.

We performed gap filling on the HiRise scaffolded assembly using Oxford Nanopore Technologies (ONT) long reads with the tool PBJelly, part of the PBSuite^47^ (version 15.8.24). PBJelly also resolved the sizes of the gaps introduced by HiRise. We used Porechop ^48^ (version 0.2.3) to adapter trim the ONT reads, resulting in 1.2 fold coverage of the genome. We set PBJelly to correct only intrascaffold gaps, and used the default minimum of 1 read spanning a gap. We reduced the number of gaps (represented in the assembly as a series of Ns) in the genome from 258,836 to 207,433 and reduced the total number of Ns in the genome from 154,284,192 to 132,359,239.

We used the genome improvement tool Pilon^49^ (version 1.22) to correct sequence errors in the gaps that were filled with the high-error ONT data. We did this by aligning the Illumina shotgun data back to the PBJelly outputted genome using bwa mem^50^ (version 0.7.7) and marking duplicates with Picard toolkit MarkDuplicates^51^ (version 1.114). We then ran Pilon with the alignment file as the “--frags” input, allowing all changes to be fixed, the PBJelly genome as the genome file, and specified the genome as diploid. We ran two iterations of alignment and consensus sequence calling.

We visually assessed this final version of the genome (PumCon1.0) by alignment to the domestic cat genome (GCA_000181335.4) using SyMap2^52^ (version 4.2) using the BLAT default alignment and synteny parameters, but with Min Dots set to 10, and Top N set to 1 (Supplementary Fig. 2). We also used the genome assessment tool BUSCO^25^ (version 2.0.1) to evaluate genome completeness based on a set of conserved single-copy orthologous genes. In the PumCon1.0 genome, 93.0% of these genes are complete, and present in a single copy only (Supplementary Table 2). Our PumCon1.0 assembly has 1.5.1QV40 unphased metric ^53^ with our confidence in the base quality of our assembly coming from prior assessments of short read data^54^.

The final genome assembly was 2,432,985,507bp in length and had an N50 of 100.53 Mb, with 178,994 gaps and 114,069,924 Ns. Ninety percent of the PumCon1.0 assembly is represented by 28 scaffolds. However, two of these scaffolds are X related (scaffold_X1 and scaffold_X2), we did not include them in further analyses. Thus 87.6% of the genome is represented on 26 autosomal scaffolds, each larger than 20 Mb. Shotgun data used for the assembly is located in the SRA under accession IDs SRR7148342-SRR7148354. The PumCon1.0 genome is available on GenBank under GCF_003327715.1.

### Genome Annotation

#### RNA-Seq library

We extracted total RNA from whole blood collected from a wild female mountain lion (SC85) from the Santa Cruz Mountains by performing a Trizol RNA extraction. We removed unwanted globin mRNA by performing the GLOBINclear protocol (Thermo Fisher). For cDNA synthesis, we used 20 ng of input RNA, a reverse transcriptase (Clonetech), a SmartSeq2 template-switch oligo^55^, and an Oligo-dT primer to enrich for poly-A+ RNA. We performed reverse transcription at 42°C for 1 hour. We treated the cDNA product with 1 µL of 1:10 dilution of RNase A (Thermo Fisher) and Lambda Exonuclease (NEB) and incubated at 37°C for 30 minutes. We amplified the cDNA for 18 cycles using KAPA Hifi Hotstart 2X readymix (KAPABiosystems) with an ISPCR primer^55^. We tagmented the amplified cDNA at 55°C for 7 minutes in a 20 µL reaction using a Tn5 enzyme to generate the RNA-Seq libraries. The Tn5 enzyme was loaded with custom oligos Tn5ME-A/R and Tn5ME-B/R^56^. We amplified the tagmented products for 13 cycles using KAPA Hifi (KAPA Biosystems) with Nextera primers. The amplified RNA-Seq library was size-selected targeting 300-800 bp using a 2% agarose E-gel (Thermo Fisher). The RNA-Seq library was then quantified using a Qubit and the size distribution was verified using an Agilent 2100 Bioanalyzer. We sent the RNA-Seq library for sequencing on a HiSeq 4000 at UC San Diego Institute for Genomic Medicine Genomics Center (2×100bp). The RNA-Seq library is publicly available on the SRA under the ID SRX4067841.

#### Annotation

The PumCon1.0 genome was annotated according to the NCBI Eukaryotic Genome Annotation Pipeline^57^ using our cDNA library and a publicly available dataset generated from a wild mountain lion from Arizona (SAMN02885420, SRX633288).

### Mountain lion resequencing data

We sequenced 11 mountain lions for this study. Ten, including the individual used for the genome assembly (SC36), were used in a panel for analysis of demographic history and population structure. One was used to assemble the X chromosome. We obtained high coverage (∼30X), whole genome sequencing data for the nine additional mountain lions that formed our panel: one from the Santa Cruz Mountains (SC29), one from Yellowstone National Park (YNP198), two from the Big Cypress National Preserve representing the canonical (last remaining authentic) Florida panther population (CYP47, CYP51), one from the Everglades National Park in Florida (EVG21), three from the Santa Monica Mountains in Southern California (SMM12, SMM22), and two from eastern Brazil (BR406, BR338). We also obtained high coverage, whole genome sequencing data for a female mountain lion from the Santa Monica Mountains for the purpose of assembling an X chromosome (SMM13).

We extracted DNA from blood samples using the Qiagen DNeasy Blood & Tissue kit using the method as used for SC36. We prepared indexed Illumina libraries following the Meyer Kircher protocol^39^. We verified DNA concentrations and ∼350-bp insert sizes by running the libraries on an Agilent 2200 TapeStation system.

We sent samples for sequencing at the National Genomics Infrastructure of SciLife in Stockholm, Sweden on an Illumina HiSeq XTen, Laboratório de Biotecnologia Animal at the Universidade de São Paulo in Brazil on an Illumina HiSeq 1500, and UC San Diego Institute for Genomic Medicine Genomics Center on an Illumina HiSeq 4000. Sample information is available on NCBI under the BioSample accessions SAMN09396574, SAMN09396570, SAMN09396599, SAMN09396617, SAMN09396621, SAMN09396547, SAMN09396626, SAMN09809203, SAMN09396627, SAMN09396580, and SAMN08662999.

We also downloaded shotgun sequencing data for the African cheetah^58^ (SRR2737512-SRR2737518) to use as the outgroup for our analyses. We selected African cheetah data with an insert size of 170 bp and obtained a final coverage of 34X when mapped to PumCon1.0.

### Variant Calling and Filtering

Prior to alignment of resequencing reads, we added the mitochondrial genome sequence for SC36 (Section Mitochondrial Genome Assemblies and Phylogeny) as a scaffold (scaffold_Mt) to the final nuclear sequence. Due to the high number of nuclear mitochondrial DNA segments (NUMTs) in felids^59^, we sought to decrease erroneous mismappings of true mitochondrial DNA in the Illumina data to the NUMTs.

We removed adapters from all mountain lion resequencing data and cheetah SRA data using SeqPrep2^42^, discarding reads shorter than 25 base pairs in length. We then aligned reads to the PumCon1.0 genome, including the mitochondrial scaffold, using bwa mem^60^ (version 0.7.12). We filtered alignments using samtools^61^ (version 1.2.1) to keep alignments with a map quality score greater than or equal to 30, remove secondary alignments, and keep reads where both reads in the read pair mapped. Within each library, we removed duplicate sequences due to PCR amplification using samtools rmdup^61^ (version 0.1.18). Realignment around insertions and deletions was performed using GATK^62^ Realigner Target Creator and Indel Realignment^62^ (version 3.5.0). We called variants in each sample using GATK HaplotypeCaller ^62^ (version 3.7.0), with a minimum base quality score of 18, and emitting all sites, including invariant positions.

We generated two sets of genotypes: one set consisting of the ten mountain lions and another set of the ten mountain lions plus the cheetah for use as the outgroup. For the mountain lion only data set, we joint genotyped all ten mountain lion samples using GATK GenotypeGVCFs^62^ (version 3.7.0) including non-variant sites. For the data set including the cheetah, we again used GATK GenotypeGVCFs to perform joint genotyping for all 11 samples, but emitted only variant sites. For both variant files, we filtered biallelic SNPs based on a number of parameters. We determined filtering thresholds by visualizing parameter distribution. We first filtered biallelic SNPs on strand odds ratio (SOR > 3.0), Fisher strand bias (FS > 60.00), quality by depth (QD < 2.00), mapping quality (MQ < 40.00 and MQRankSum < −10.00), read position (−8.00 < ReadPosRankSum > −8.00), and excess heterozygosity in accordance with Hardy-Weinberg equilibrium (ExcessHet > 10.0) using GATK VariantFiltration and SelectVariants^62^. For the mountain lion only variant file, we removed variants with a cumulative depth greater than 1500 (DP > 1500), while we removed variants with a cumulative depth greater than 1650 for the mountain lion and cheetah variant file (DP > 1650) using GATK VariantFiltration and SelectVariants^62^. We also removed singleton variants where only one individual carried one copy of a different allele (vcftools--mac 2), sites where any individual had a depth less than 10 (vcftools --minDP 10.0), and sites where any one of the ten mountain lions did not have a base called (vcftools --max-missing-count 0)^63^. We only used autosomal scaffolds for further analyses, removing mitochondrial or X chromosome related scaffolds (scaffold Mt, X1, X2, X3, 869, 1862). Scaffolds 869 and 1862 were removed from the variants due to syntenic mappings between them and the X chromosome of FelisCatus9.0 in SyMap^52^. We performed linkage disequilibrium filtering on the both variant files using PLINK^64^ (version 1.90b4.4) with the command “--indep-pairwise 100 10 0.25”. The final mountain lion only variant file contained 166,037 sites. The mountain lion and cheetah variant file contained 557,741 sites. The larger number of variants in the mountain lion and cheetah file is due to the high numbers of sites where the cheetah carries two of the alternate allele, while all mountain lions carry the reference allele.

Using the non-LD filtered SNP calls from the mountain lion only data set, we generated a fasta file for each sample using GATK FastaAlternateReferenceMaker^62^ (version 3.7.0), with heterozygous positions represented by IUPAC codes. Failed SNP sites (see filters above) and failed individual genotypes (RGQ<20, depth<10) were masked to Ns using the *maskfasta* function in bedtools^65^ (version 2.25.0).

### Mitochondrial Genome Assemblies and Phylogeny

We used Unicycler^66^ (version 0.4.4) to assemble an initial mitochondrial sequence for SC36. We first identified mitochondrial mapping reads to decrease our initial input dataset. To do this, we mapped adapter-trimmed Illumina and ONT reads to a publicly available mountain lion reference mitochondrial sequence (KP202261.1)^67^ using bwa mem^50^ (version 0.7.12) and bwa mem ont2d, respectively. We took the reads in the Illumina and ONT alignment files that mapped to the reference mitochondrial sequence and converted them into fastq format using bedtools^65^ bamToFastq. We used the Illumina and ONT mitochondrial mapped readsets as input into Unicycler as long and unpaired reads, respectively. The assembly created with Unicycler was 17,065bp in length, circular, and had a depth of 490X. To validate this assembly, we used the Unicycler output as the reference sequence in an assembly using mapping iterative assembler (mia)^68^ (version 1.0) using roughly 20 million randomly selected adapter-trimmed Illumina shotgun reads (command: mia -i -c -C -F -k 13). The resulting mia assembly was identical in sequence to the inputted Unicycler assembly, and thus was the final mitochondrial sequence for SC36.

We used adapter-trimmed Illumina shotgun data to assemble the mitochondrial genome sequences of the remaining nine mountain lions with mia, using the SC36 mitochondrial sequence as the reference. The coverages of these mitochondrial assemblies ranged from 35X to 138X. For each of the nine assemblies, we filtered using a consensus threshold of 90% and required 10X coverage per site. A site which did not meet this requirement were changed to an ‘N’, resulting in between 2 and 86 Ns per assembly. We removed two repetitive sequences located in the control region, RS2 (sites 16,511-16,861) and RS3 (sites 279-657), from each of the mountain lions and cheetah mitochondrial sequences. These regions are known to have highly variable numbers of repeats^69^–^72^, and were difficult to accurately resolve using short read data. We annotated the mitochondrial genomes using MITOS^73^. The annotated mitochondrial assemblies for the ten mountain lion are available on GenBank under accession numbers MH807447, MH814703-MH814707, and MH818219-MH818222.

We ran PartitionFinder^74^ (version 1.1.1) and jModelTest2^75^ (version 2.1.6) to determine the partitions and best substitution model. We kept the mitochondria in one partition due to the small number of substitutions seen in the mountain lions, and used a GTR+GAMMA substitution model for tree building.

We used muscle^76^ (version 3.8.31) to align our ten assembled mountain lion mitochondrial genomes, the publicly available mountain lion reference mitochondrial sequence (downloaded from GenBank, KP202261.1), and a cheetah mitochondrial sequence (downloaded from GenBank, accession KP202271.1) as the outgroup. We used RaxML^77^ (version 8.2.4) to produce a maximum likelihood phylogeny, with a GTR+GAMMA evolutionary model, running one hundred bootstrap replicates. We also ran tree inference including the RS2 and/or RS3 repetitive regions, and saw little change in the bootstrap support values and no change to the topology of the tree.

We estimated divergence times for South and North American mountain lions using a prior composite estimate of the feline mitochondrial divergence rate of 1.15%bp/Myr^10,70^. Branch lengths were taken from a phylogeny that did not include RS2 or RS3, and were used to calculate divergence dates using the formula σ = 2λT, where σ is percent divergence between a pair of sequences, λ is the rate of mitochondrial divergence [1.15%bp/Myr], and T is time^70^. All mountain lion mitochondrial sequences diverged 278,000 ± 5,639 years ago, while North American mitochondria diverged from South America 201,000 ± 1,952 years ago, and divergence within North America occurred 21,000 ± 10,412 years ago.

### Demographic History

We used the pairwise sequentially Markovian coalescent (PSMC) model^78^ to estimate the historical effective population size of each mountain lion in this study. The input was a realigned alignment file of 26 autosomal scaffolds larger than 20 Mb, excluding sex chromosome associated scaffolds (Section: Variant calling and filtering). We filtered the alignment file for each mountain lion to include sites which had from one third the average coverage for that mountain lion to twice the average coverage for that mountain lion. We used a generation time of 5 years and a per generation mutation rate of 0.1e-8, based on previous estimates of the feline mutation rate^79^. We performed one hundred replicate bootstraps for each individual per the software instructions (Supplementary Fig. 4).

We also ran the PSMC tool on regions of the alignment file which were identified as outbred based on our ROH hidden Markov model (Section Runs of Homozygosity). We masked regions of homozygosity for each of the mountain lions, and re-ran the PSMC analysis to compare the demographic estimates. We saw no considerable difference between the two sets of results (Supplementary Fig. 5).

### Population Structure

We used SmartPCA from the EIGENSOFT^80^ (version 6.1.4) package to run principal component analysis on the LD-filtered variant file for the ten mountain lions. We converted the variant file into eigenstrat format using PGDSpider2^81^ (version 2.1.0.0) for input into SmartPCA.

We constructed a tree to show population splits using Treemix^82^, both with and without the admixed sample EVG21. We used the LD-filtered variant file with the ten mountain lions and the cheetah as input, and ran Treemix using a SNP window size of 5,000. For the dataset including EVG21, the tree with the highest log likelihood predicted 1 migration, and explained 99.91% of the variation. In the dataset without EVG21, the best model predicted no migrations and explained 99.91% of the variation.

We used the software STRUCTURE^83^ (version 2.3.4) to infer the population structure of the mountain lions. We converted the LD filtered variant file into the input format using PGDSpider2^81^ (version 2.1.0.0). We ran 10 replicates of STRUCTURE for values from one to ten for the number of populations (K) using a degree of admixture (alpha) value of 1 for each K and an admixture model, 10,000 burns ins, and running 10,000 MCMC repetitions. We identified the best K based on the highest delta K value^84^. For our dataset, K=3 had the largest delta K value (Supplementary Fig. 7). We used the software CLUster Matching and Permutation Program (CLUMPP)^85^ (version 1.1.2) under the greedy algorithm to align the ten replicates for each K into one representative mean output matrix for plotting.

### Genome-wide Heterozygosity Estimates

We calculated the average coverage of each mountain lion using samtool depth on the realigned alignment file, removing scaffolds for the mitochondria and X chromosome (scaffolds Mt, X1, X2, X3, 869, 1862). We calculated genome-wide heterozygosity by generating a pileup file from map quality and base quality filtered data (samtools mpileup -q 30 -Q 30) at all sites in the genome with exactly the average coverage depth for that individual. We created a histogram by binning the sites based on the number of reads representing the reference allele. We visually classified which bins were designated as homozygous or heterozygous. We calculated the genome wide heterozygosity by summing all heterozygous bins and dividing by the total number of genome wide sites used in the analysis. We validated our heterozygosity estimates using the --het flag in PLINK^64^ (version 1.90b4.4) (Supplementary Table 3). Additionally, we examined sliding window heterozygosity using a custom script that counts heterozygous positions in the IUPAC coded fasta files, estimating average heterozygosity in 100-kb windows. We used the outputted counts of the number of heterozygous positions and the number of positions with a genotype call to obtain another estimate of average genome-wide heterozygosity.

### Runs of Homozygosity

We used a hidden Markov model (HMM) to identify ROH^86^. We first estimated HMM model parameters from the data (Supplementary Table 4). For each of the eight male mountain lions, we used the sample’s heterozygosity estimate from the two large X scaffolds as an estimate of the genotyping error rate. For the two female mountain lions, we used the estimate from another sample that was geographically close by and had similar sequencing coverage. For each sample, we then estimated the rate of heterozygosity in outbred regions by visually selecting clearly outbred regions and determining the mean heterozygosity across those regions. We determined a single transition rate parameter (t=1e-50) by running different transition parameters and visually inspecting the results. We used these parameters and the filtered fasta files with IUPAC codes as input into the ROH HMM program that identifies transitions between inbred and outbred regions of the genome. For downstream analyses, we used only the 26 largest autosomal scaffolds and discarded ROH less than 2 Mb. We converted the ROH tract lengths to generations using an estimated average recombination rate in the domestic cat of 1.1 cM/Mb^87^ and the equation g = 100/(2rL), where g is the time in generations, r is the recombination rate, and L is the length of the ROH tract in Mb^38^.

We also used the sliding window approach in PLINK^88^ (version 1.90b4.4) to identify ROH for comparison with our ROH HMM. We used the quality filtered, non-LD filtered mountain lion vcf as input. We relaxed the parameters (--homozyg-window-het 20 --homozyg-window-missing 20 --homozyg-window-threshold 0.02 --homozyg-het 750 --homozyg-kb 500) to prevent sequencing errors from breaking up homozygous tracts. Even with the relaxed parameters, PLINK still tended to break up long tracts (Supplementary Fig. 10). Since accurate estimates of tract lengths were key to our inbreeding analysis, we used the ROH called by the custom HMM program for further analyses.

We used Ancestry_HMM^89^ to classify ancestry in EVG21 into three types: homozygous Central/South American, homozygous Floridian, and heterozygous. We used the two Brazil samples (BR338 and BR406) as proxies for Central/South American ancestry, and the two Big Cypress samples (CYP47 and CYP51) as proxies for Floridian ancestry. Because small sample sizes preclude accurate estimates of LD and because the program is sensitive to sites in strong LD, we pruned all ancestry informative markers within 250 kb of another site. The ancestry types approximately followed a Hardy-Weinberg distribution, where homozygous Brazilian ancestry was 22% of the genome, homozygous Floridian ancestry was 28% of the genome, and heterozygous ancestry was 50% of the genome. We used the ROH calls from our ROH HMM program to identify ROH greater than 2 Mb in each ancestry type. We did this by using the *intersect* function in bedtools^65^ (version 2.25.0) with the parameters “-e -f 0.90” such that 90% of a ROH was one ancestry type as designated by our ancestry HMM. We saw no ROH greater than 2 Mb that were classified as heterozygous, thus admixture effectively breaks down long tracts of homozygosity (Supplementary Fig. 12). We tested for significance differences between the number of ROH for each ancestry type using a poisson distribution. We tested for significance in the number of ROH identified per ancestry type using an exact test of the ratio with a Poisson distribution. We found that more ROH were of Floridian ancestry relative to heterozygous ancestry with a p-value of 7.276e-12, and more ROH were of Central/South American ancestry than of heterozygous ancestry with a p-value of 1.526e-05. Additionally, more ROH were of Floridian ancestry than Central/South American ancestry, with a p-value of 0.006456.

We next estimated the proportion of the ROH that are shared between pairs of mountain lions. First, we used the *intersect* function in bedtools^65^ (version 2.25.0) to find genomic regions where ROH overlap between pairs of samples. Then we used a custom script to generate a hybrid diploid fasta file from the IUPAC coded fasta files for each pair of samples. The script first makes a pseudo-haplodized sequence for each sample by randomly selecting one of the two bases at heterozygous sites. The script then uses a pair of pseudo-haplodized sequences to generate a hybrid diploid fasta file by using IUPAC codes to represent differing bases between the two samples, while shared bases between the two samples are represented by the base itself. We ran the ROH HMM program on the generated fasta file, using the same transition rate parameter as above (t=1e-50) and the outbred heterozygosity rate and genotyping error rate averaged across all ten mountain lions (h=0.00135, e=0.0000765). These ROH indicate where two individuals share regions of the genome IBD. To determine if ROH that overlap between two samples were also IBD, we used the *intersect* function in bedtools^65^ (version 2.25.0) to find the intersection between the ROH that overlap between the two samples and the ROH generated from the hybrid diploid fasta file. Finally, from these outputs, for each pair of mountain lions we calculated the proportion of the genome that occurs in ROH that are IBD.

## Supporting information

## Acknowledgements

We thank the Wilmers Lab for helping to collect samples, as well as R. Miotto, E. Amorim, J. May, CENAP/ICMBio/Brazil and AMC/Brazil for access to biological samples; S. Webber from the Paleogenomics lab, and C. Scelfo-Dalbey for assistance in generating data. The authors would like to acknowledge support from Science for Life Laboratory, the National Genomics Infrastructure, and UPPMAX for providing assistance in massive parallel sequencing and computational infrastructure. Sequencing was also performed by the Laboratório de Biotecnologia Animal at the Universidade de São Paulo in Brazil, UC San Diego Institute for Genomic Medicine Genomics Center, UC Berkeley Vincent J. Coates Genomics Sequencing Laboratory, and UC Santa Cruz Ancient and Degraded Processing Center.

## Author contributions

BS, REG, and CCW conceived and designed the study. CCW, DPO, DRS, EE, HVF, LD, LLC, PMSV, RKW, SJO, and WJ provided samples or data for this work. AB, BO, HJM, and PMSV performed laboratory work. BS, REG, and RCD supervised the analysis. RCD, NFS, MAS, and JAC analyzed the data. BS, RDG, RCD, NFS, MAS, CCW, WJ, SJO, RKW, and EE interpreted the results. BS, NFS, and MAS wrote the manuscript. All authors edited the manuscript.

## Author information

Funding for the mountain lion genome was provided by a grant to CCW from the Gordon and Betty Moore Foundation. BS, MAS, NFS, and RKW were funded by a grant from the University of California Office of the President. LD was funded by Formas grant nr 2015-676. DRS was funded in part by Yellowstone Forever. BS and REG were funded in part from NSF DEB-1754551. EE, LLC and HVF were supported by funds from CNPq/Brazil and INCT-EECBio/Brazil.

## Additional Information

### Data Availability

Shotgun data used for the assembly have been deposited in the SRA under accession IDs SRR7148342-SRR7148354. The PumCon1.0 genome is available on GenBank under GCF_003327715.1. The RNA-Seq library is publicly available on the SRA under the ID SRX4067841. Reads for the panel of mountain lions have been deposited in the SRA under the accession IDs SRR7639695-6, SRR7542886-8, SRR7660678-9, SRR7664677-8, SRR7956993-4, SRR7610940-1, SRR7661934-5, SRR7690239-40, SRR7543017-8, SRR7537344-5, and SRR7148342-54. The annotated mitochondrial assemblies for the ten mountain lion are available on GenBank under accession numbers MH807447, MH814703-MH814707, and MH818219-MH818222.

