## Supplementary material for "Mountain lion genomes provide insights into genetic rescue of inbred populations"

### Supplementary information

**Supplementary Table 1: Genome assembly metrics.** The metrics of the different versions of the mountain lion genome. The final column represents the final assembly, PumCon1.0. We saw a marked improvement in the N50 as a result of scaffolding with HiRise. Gap filling with PB Jelly notably decreased the numbers of Ns and strings of Ns in the genome assembly. Due to gap filling and the creation of previously missing regions of the genomes, more Illumina reads mapped to the genome assembly. Due to the error frequency of ONT reads, the correction of the gap filled region sequence with iterative rounds of Pilon using the Illumina data also increased the number of reads that mapped to the genome. Asterisk denotes that the Meraculous assembly did not include the final versions of the X chromosome scaffolds.

| | Meraculous | HiRise | PBJelly | Pilon iteration 1 | Pilon iteration 2 (final) |
|---|---|---|---|---|---|
| Assembly step | Shotgun assembly | Scaffolding | Gap Filling | Error correcting | Error correcting |
| Input data | Illumina shotgun reads | Chicago & Hi-C libraries | ONT reads | Illumina shotgun reads | Illumina shotgun reads |
| Genome length (bp) | 2,181,316,782 | 2,293,137,739 | 2,433,777,904 | 2,433,231,347 | 2,432,985,507 |
| N50 scaffold | 36.6kb | 103.78 Mb | 100.51 Mb | 100.54 Mb | 100.53 Mb |
| L50 scaffold | 17,135 | 7 | 8 | 8 | 8 |
| # of gaps | 124,710 | 258,836 | 207,433 | 184,611 | 178,994 |
| # of Ns | 26,631,327 | 154,284,192 | 132,359,239 | 119,328,697 | 114,069,924 |
| % of Ns in genome | 1.22% | 6.73% | 5.44% | 4.90% | 4.69% |
| # Illumina shotgun reads mapping (samtools view -q 30 -c) | 958,095,130* | 979,540,843 | 982,339,501 | 985,819,801 | 987,305,346 |

**Supplementary Table 2: Benchmarking Universal Single-Copy Orthologs (BUSCO) gene completeness score**. The results of running BUSCO[1] on the PumCon1.0 genome using the human gene set (n=4104).

| | |
|---|---|
| Complete BUSCOs | 3832 (93.4%) |
| Complete and single-copy BUSCOs | 3815 (93.0%) |
| Complete and duplicated BUSCOs | 17 (0.4%) |
| Fragmented BUSCOs | 141 (3.4%) |
| Missing BUSCOs | 131 (3.2%) |

**Supplementary Table 3: Details for the panel of mountain lions used in this study.** *Note: SMM13 was used solely for the X chromosome scaffold assembly, and thus further metrics (i.e. heterozygosity) were not calculated in this study.

| Mountain lion | Population | Gender | Cover age | Heterozygosity (pileup method) | Heterozygosity (PLINK 26 autosomal scaffolds) | Proportion of genome in an ROH | Alternate ID names | SRA accession IDs |
|---|---|---|---|---|---|---|---|---|
| BR338 | Minas Gerais state, Brazil (BR) | male | 48x | 0.00155 | 0.00101 | 0.06736 | bPco338 | SRR7639695-6 |
| BR406 | São Paulo state, Brazil (BR) | male | 27x | 0.00166 | 0.00103 | 0.03970 | D406 | SRR7542886-8 |
| EVG21 | Everglades National Park (EVG) | female | 51x | 0.00121 | 0.00091 | 0.34221 | FP021, Pco-007 5 | SRR7660678-9 |
| CYP47 | Big Cypress National Preserve (CYP) | male | 43x | 0.00033 | 0.00032 | 0.58485 | FP047, Pco-042 3 | SRR7664677-8 |
| CYP51 | Big Cypress National Preserve (CYP) | male | 55x | 0.00034 | 0.00034 | 0.56168 | FP051, Pco-042 8 | SRR7956993-4 |
| YNP198 | Yellowstone National Park (YNP) | male | 40x | 0.00090 | 0.00077 | 0.15873 | M198 | SRR7610940-1 |
| SMM12 | Santa Monica Mountains (SMM) | male | 46x | 0.00079 | 0.00073 | 0.18884 | P12 | SRR7661934-5 |
| SMM13* | Santa Monica Mountains (SMM) | female | 40x | NA | NA | NA | P13 | SRR7690239-40 |
| SMM22 | Santa Monica Mountains (SMM) | male | 34x | 0.00059 | 0.00049 | 0.417637 | P22 | SRR7543017-8 |
| SC29 | Santa Cruz Mountains (SC) | female | 35x | 0.00062 | 0.00058 | 0.32707 | 29F | SRR7537344-5 |
| SC36 | Santa Cruz Mountains (SC) | male | 47x | 0.00049 | 0.00050 | 0.33987 | 36M | SRR7148342-54 |

**Supplementary Table 4: ROH HMM parameters.** Parameters used as input into the HMM ROH script for each mountain lion.

| Sample | Genotyping error rate | Outbred heterozygosity |
|---|---|---|
| BR338 | 0.000146 | 0.0019 |
| BR406 | 0.000054 | 0.0018 |
| EVG21 | 0.000071 | 0.0019 |
| CYP47 | 0.000078 | 0.0011 |
| CYP51 | 0.000071 | 0.0010 |
| SC29 | 0.000078 | 0.0012 |
| SC36 | 0.000027 | 0.0010 |
| SMM12 | 0.000075 | 0.0012 |
| SMM22 | 0.000078 | 0.0012 |
| YNP198 | 0.000087 | 0.0012 |

**Supplementary Table 5: ROH pairwise IBD values.** Percent of the genome in an IBD ROH between pairs of mountain lions as shown in Fig. 4D.

| | BR338 | BR406 | EVG21 | CYP47 | CYP51 | SC29 | SC36 | SMM12 | SMM22 | YNP198 |
|---|---|---|---|---|---|---|---|---|---|---|
| BR338 | 6.74% | | | | | | | | | |
| BR406 | 0.00% | 3.97% | | | | | | | | |
| EVG21 | 0.00% | 0.00% | 34.22% | | | | | | | |
| CYP47 | 0.00% | 0.00% | 8.72% | 58.49% | | | | | | |
| CYP51 | 0.00% | 0.00% | 6.85% | 35.87% | 56.17% | | | | | |
| SC29 | 0.00% | 0.00% | 0.19% | 1.17% | 1.07% | 32.71% | | | | |
| SC36 | 0.00% | 0.00% | 0.09% | 0.70% | 0.70% | 11.55% | 33.99% | | | |
| SMM12 | 0.00% | 0.00% | 0.00% | 0.60% | 0.48% | 2.96% | 2.72% | 18.88% | | |
| SMM22 | 0.00% | 0.00% | 0.09% | 0.92% | 0.76% | 5.10% | 4.16% | 4.39% | 41.76% | |
| YNP198 | 0.00% | 0.00% | 0.18% | 0.30% | 0.22% | 0.56% | 0.16% | 0.67% | 0.56% | 15.87% |

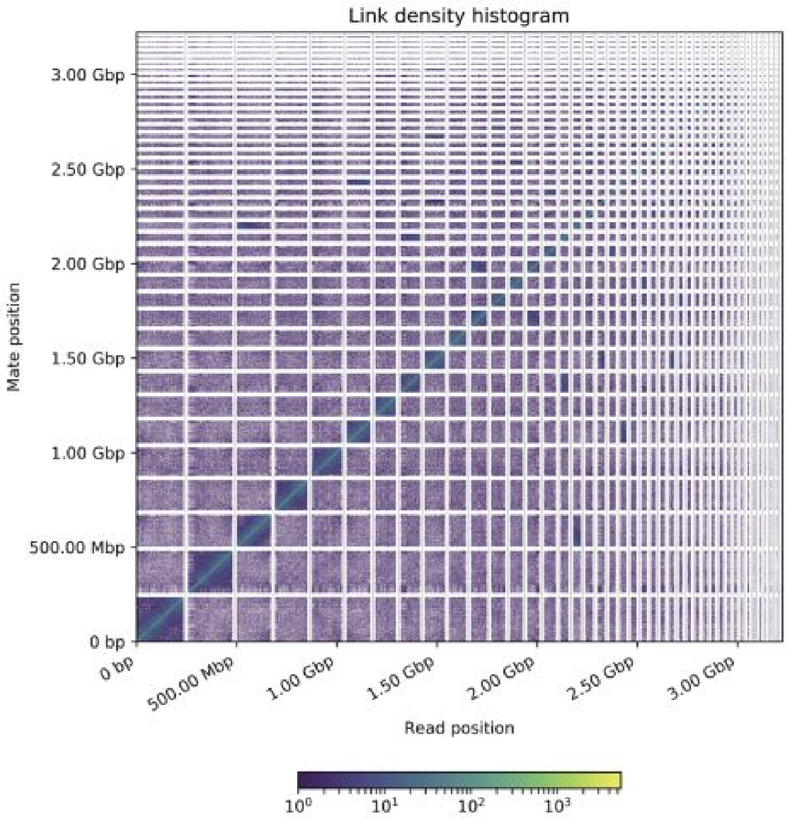


**Supplementary Figure 1: Linkage map of HiRise genome assembly for SC36.** The x and y axes mark the mapping positions of the first and second read in a read pair of the Hi-C library respectively, grouped into bins which represent scaffolds. Each square is colored according to the number of read pairs within the bin. Scaffolds less than 1 Mb in length are not shown.

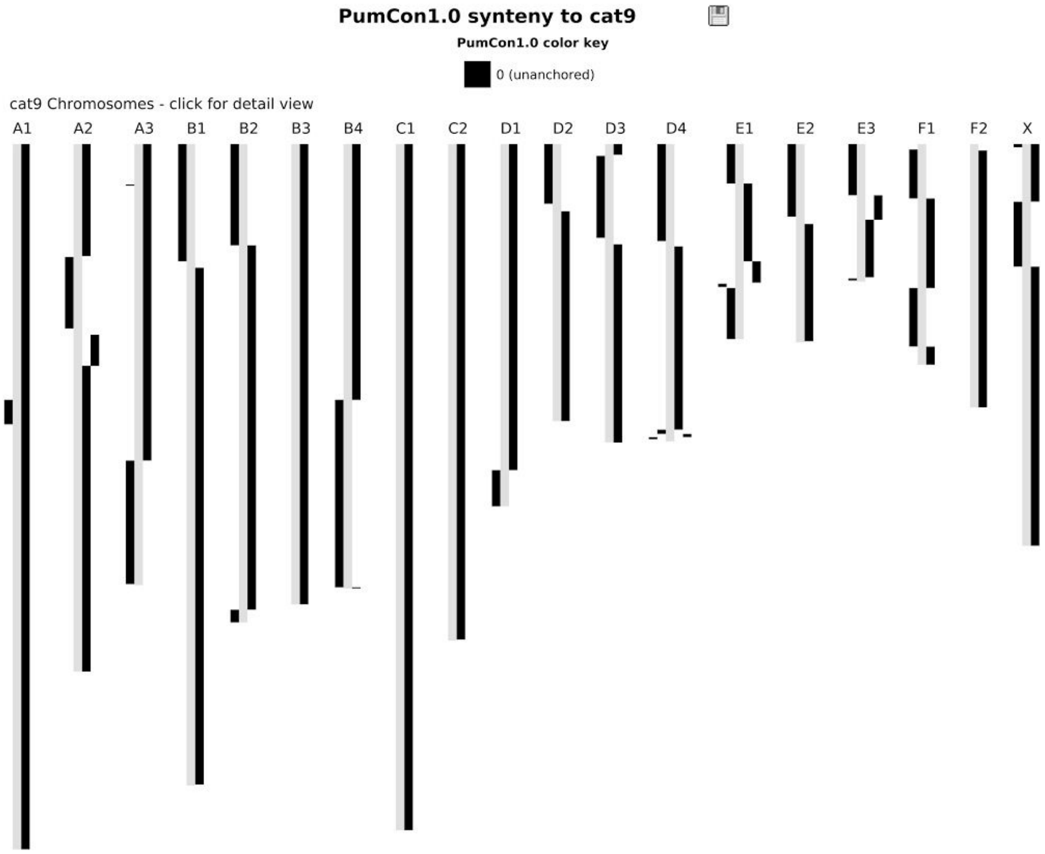


**Supplementary Figure 2: Synteny between PumCon1.0 and Felis catus 9.0 genome.** Alignment of syntenic regions of the Felis catus 9.0 (domestic cat, gray) genome assembly with our PumCon1.0 mountain lion assembly (black) using SyMap2[2]. Four of the scaffolds in our assembly mapped to the entirety of a chromosome in the cat genome.

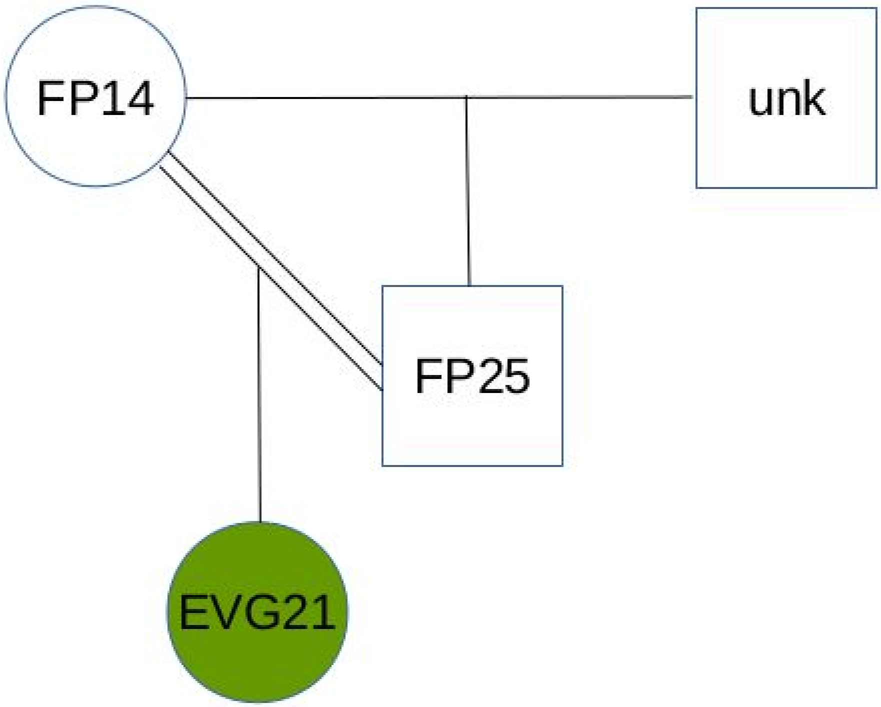


**Supplementary Figure 3: Pedigree of EVG21.** Pedigree of the inbred and admixed Florida panther sequenced from Everglades National Park[3]. All panthers in the pedigree are of Everglades ancestry. The Central American admixture into this population occurred approximately 6-9 generations prior to these individuals[3]. Note: EVG21 is referred to as FP021 in the original dataset[3].

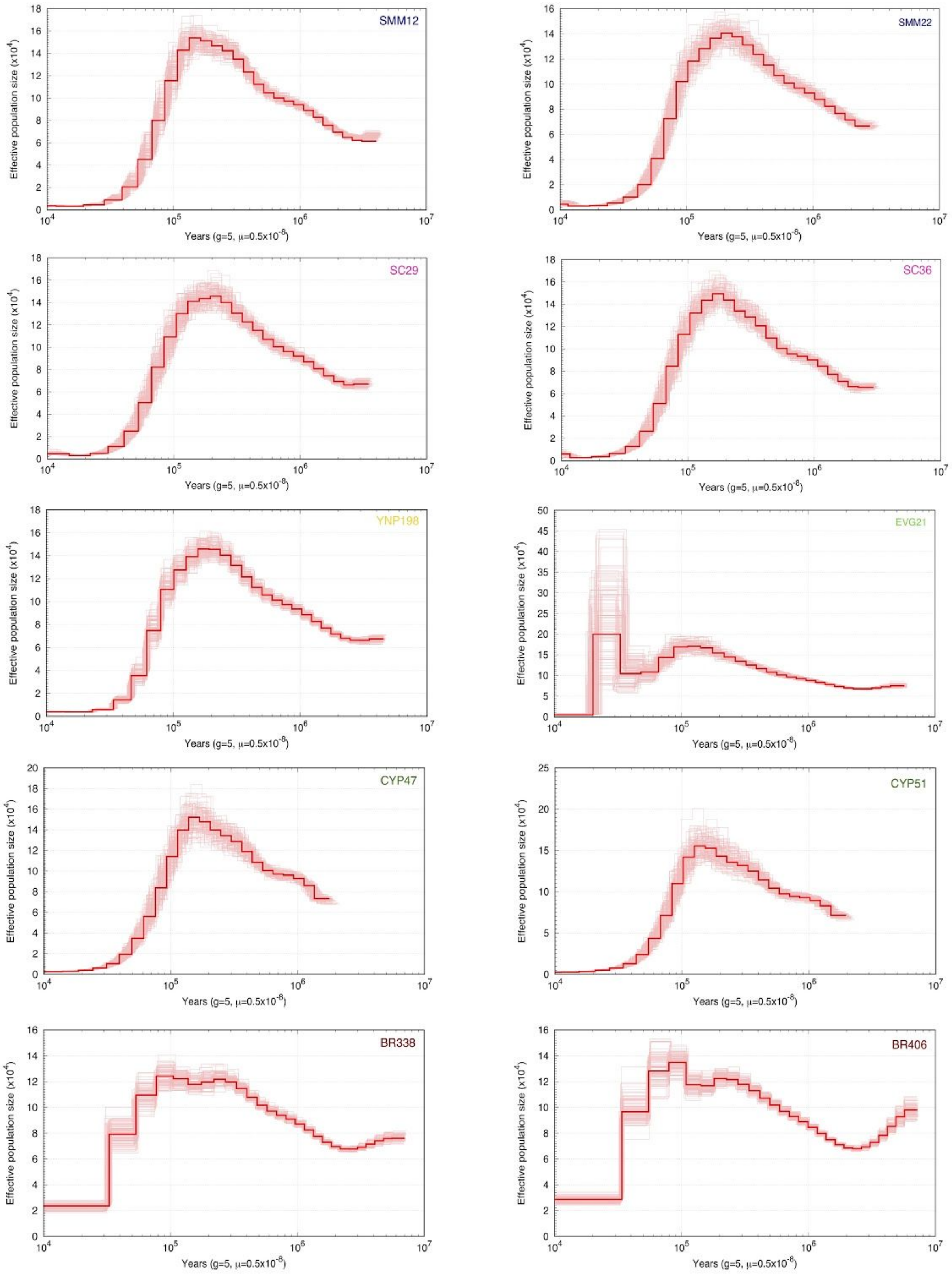


**Supplementary Figure 4: Bootstrap replicate PSMC plots for ten mountain lions.** We ran one hundred bootstrap replicates for each of the mountain lions using the PSMC model[4]. Generation time is 5 years, and per generation mutation rate is 0.5e-8[5].

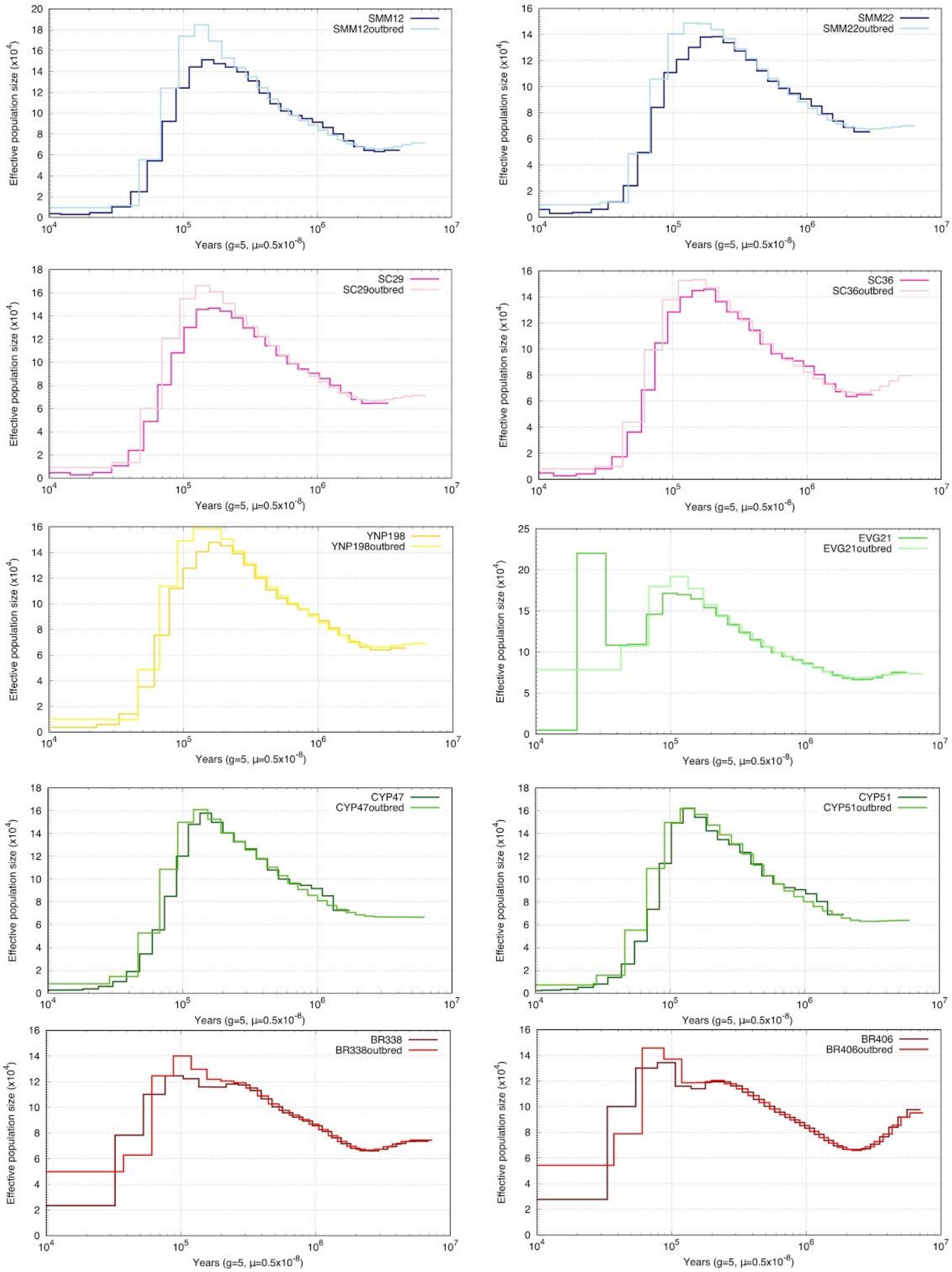


**Supplementary Figure 5: PSMC plots for outbred (non-ROH) and entire genomes of the ten mountain lions.** We saw no significant difference between the two models[4]. Generation time is 5 years, and per generation mutation rate is 0.5e-8[5].

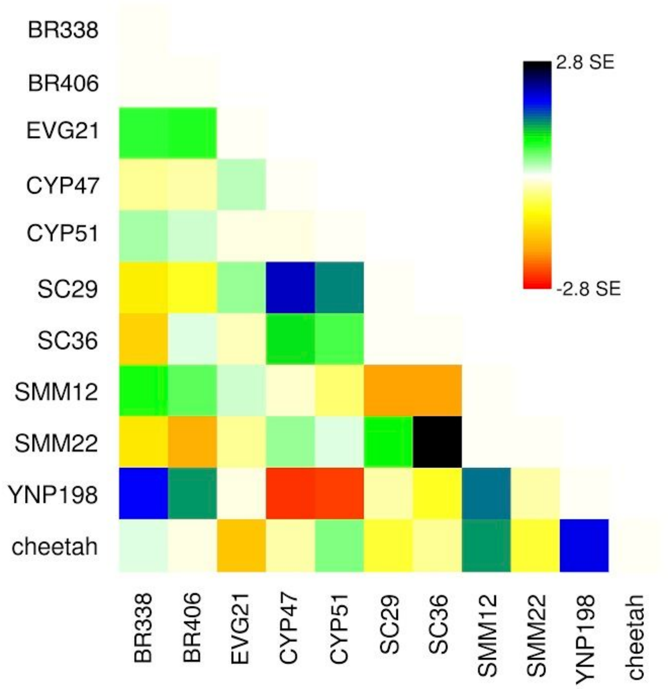


**Supplementary Figure 6: TreeMix residual fit of the model.** TreeMix[6] run on LD filtered variant file using ten mountain lions and African cheetah, 1 migration, and k =5000, where 99.91% of the variation is explained.

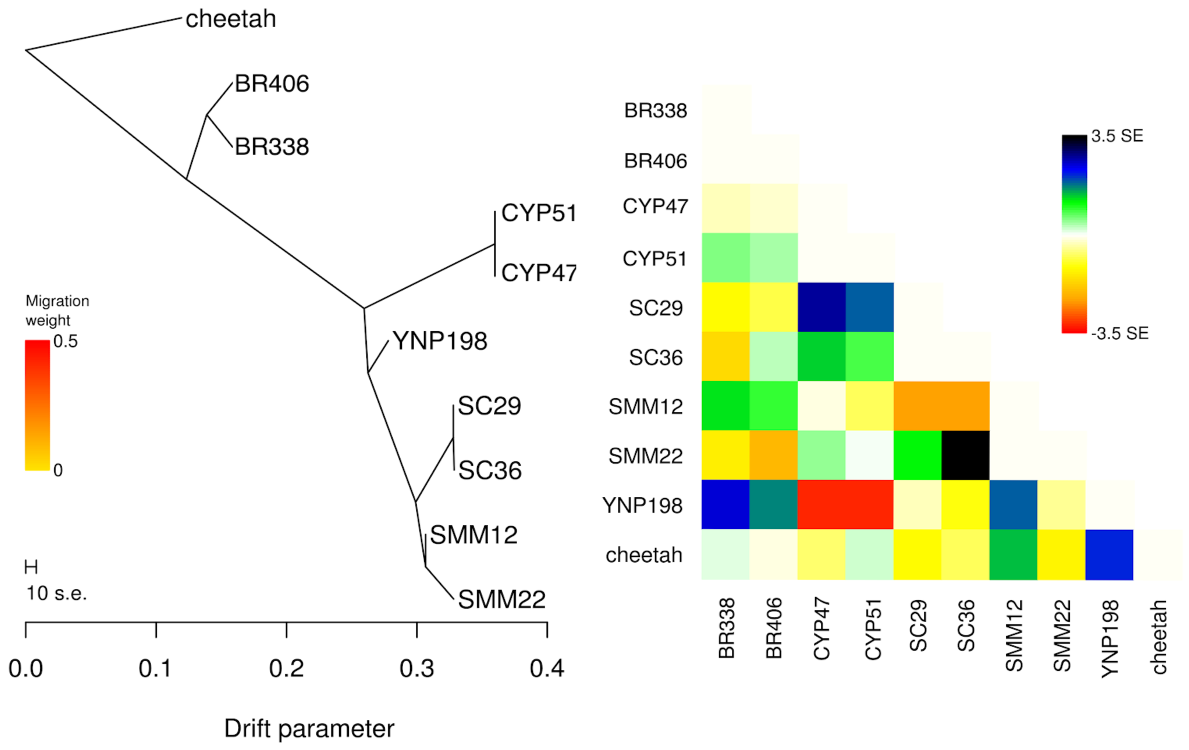


**Supplementary Figure 7: TreeMix without EVG21**. Best result of TreeMix[6] run on LD filtered variant file using nine mountain lions and African cheetah, no migrations, and k= 5000, where 99.91% of the variation is explained.

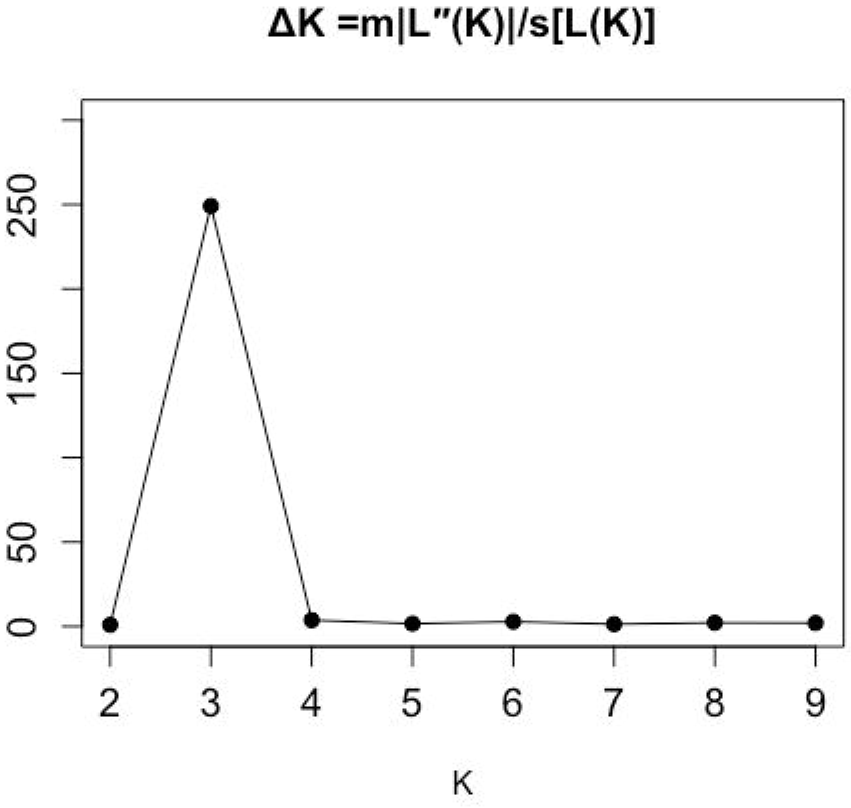


**Supplementary Figure 8: Selection of best K in STRUCTURE.** Using delta K[7], the rate of change in the log probability of data between successive K values, we identified three as the best K for our panel.

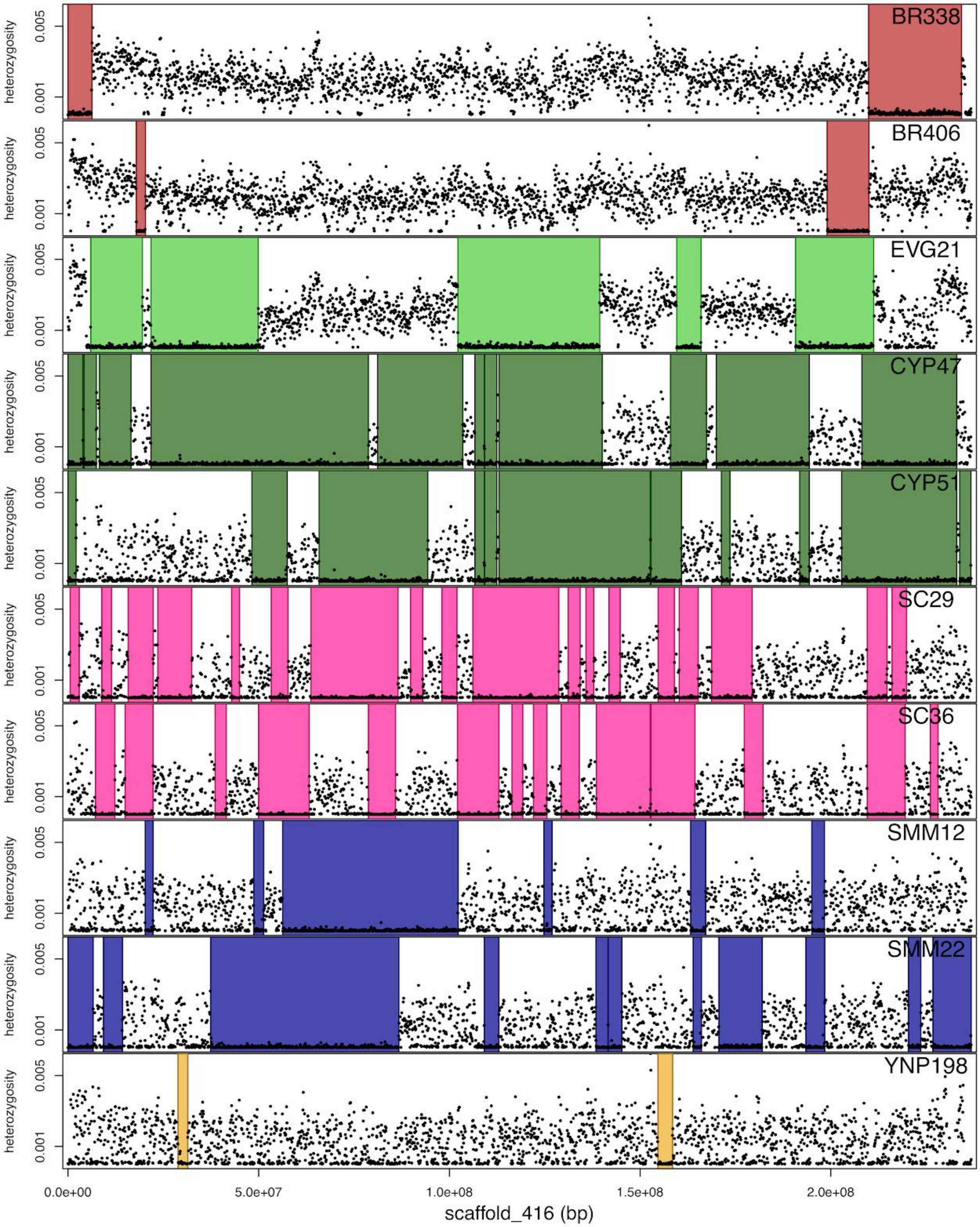


**Supplementary Figure 9: ROH for all ten mountain lions as called by our ROH HMM.** Black dots represent average heterozygosity in 100 kb windows; colored regions represent blocks called as ROH. Scaffold_416 represents the largest scaffold in the genome assembly at 236.8 Mb.

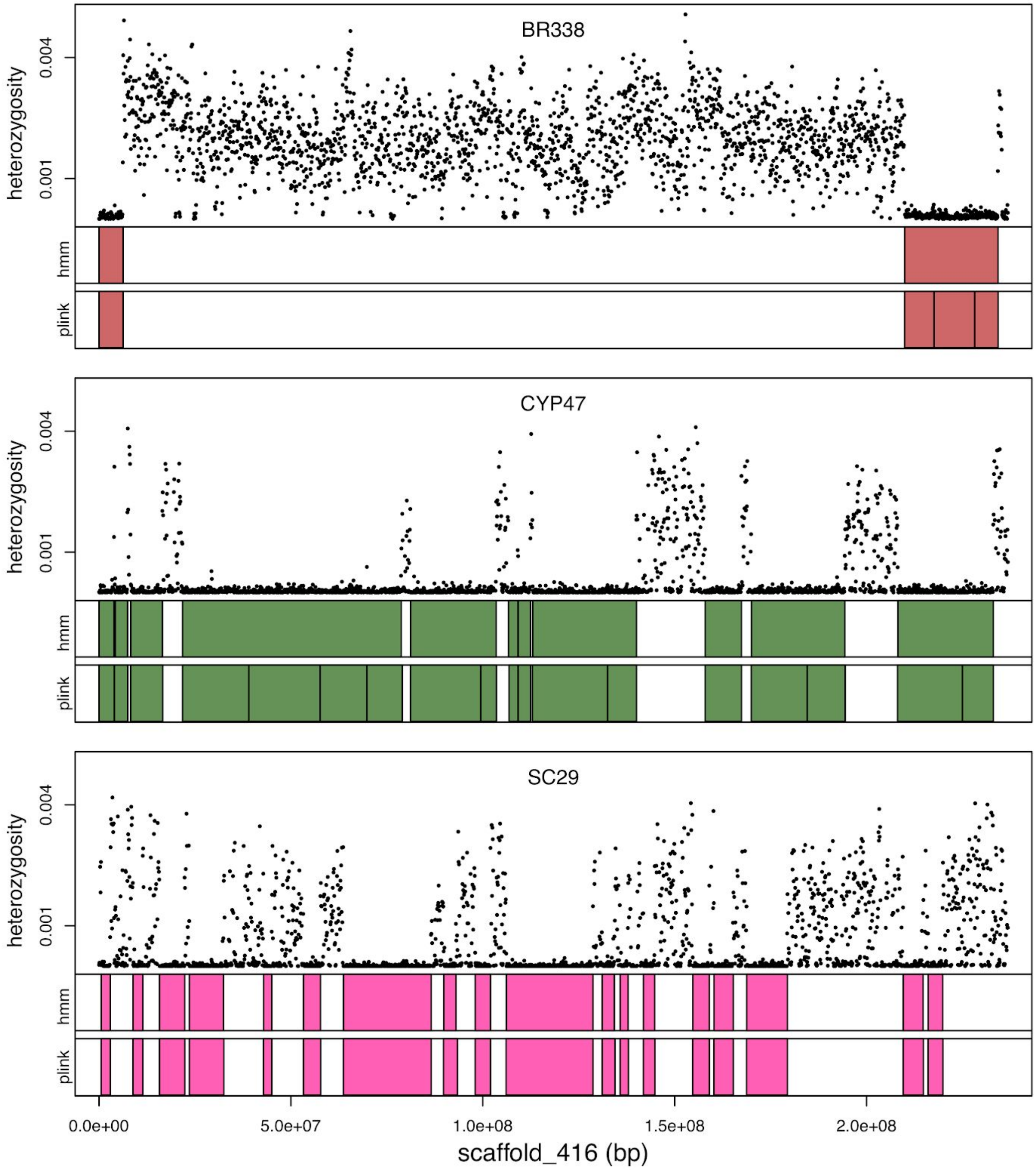


**Supplementary Figure 10: ROH calls using two different methods.** Top panel for each sample shows heterozygosity in 100 kbp windows. Bottom panels for each sample shows colored boxes indicating ROH called using our ROH HMM (top), and PLINK (bottom). PLINK tended to break up long tracts of ROH. Given that we were interested in the distribution of ROH lengths, we decided to use our HMM for ROH calls for further analyses.

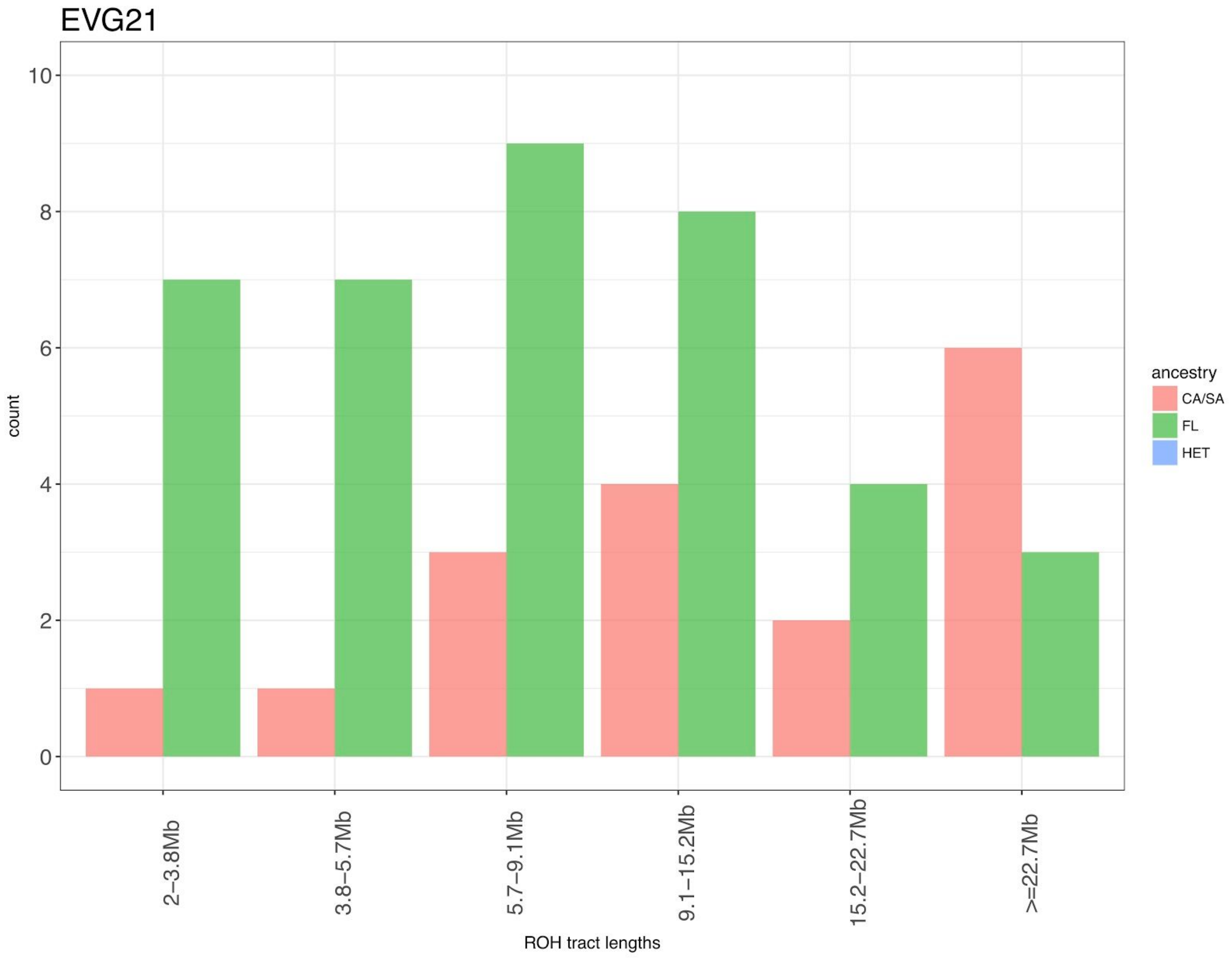


**Supplementary Figure 11: Ancestry of ROH in EVG21 genome.** We assigned ancestry to the haploid fasta sequence of EVG21 using an ancestry HMM, subdividing the ancestry into three types: homozygous Central/South American ancestry, heterozygous ancestry, or homozygous Floridian ancestry. We identified which ancestry type each ROH > 2 Mb. We observed no ROH greater than 2 Mb in length in heterozygous ancestry regions for EVG21. Admixture effectively rescued the long tracts of homozygosity. We still saw ROH of greater than 2 Mb for both homozygous Central/South American ancestry and homozygous Floridian ancestry, both at a significant occurrence relative to the occurrence of heterozygous ancestry ROH.
